# Structure of the PfRCR complex which bridges the malaria parasite and erythrocyte during invasion

**DOI:** 10.1101/2023.01.30.526221

**Authors:** Brendan Farrell, Nawsad Alam, Melissa N. Hart, Abhishek Jamwal, Robert J. Ragotte, Hannah Walters-Morgan, Simon J. Draper, Ellen Knuepfer, Matthew K. Higgins

**Affiliations:** Department of Biochemistry, Dorothy Crowfoot Hodgkin Building, University of Oxford, South Parks Rd, Oxford, OX1 3QU; Kavli Institute for Nanoscience Discovery, Dorothy Crowfoot Hodgkin Building, University of Oxford, South Parks Rd, Oxford, OX1 3QU; The Royal Veterinary College, Hawkshead Lane, North Mymms, Hatfield, Hertfordshire, AL9 7TA

**Author notes:** These authors contributed equally (Nawsad Alam, Melissa N. Hart).

## Abstract

The symptoms of malaria occur during the blood stage of infection, when parasites invade and replicate within human erythrocytes. The five-component PfPCRCR complex, containing PfRH5, PfCyRPA, PfRIPR, PfCSS and PfPTRAMP, is essential for erythrocyte invasion by the deadliest human malaria parasite, Plasmodium falciparum. Invasion can be prevented by antibodies or nanobodies against each of these five conserved proteins, making them the leading blood stage malaria vaccine candidates. However, little is known about the molecular mechanism by which PfPCRCR functions during invasion. Here we present the structure of the PfRCR complex, containing PfRH5, PfCyRPA and PfRIPR, determined by cryogenic-electron microscopy. This reveals that PfRIPR consists of an ordered multi-domain core flexibly linked to an elongated tail. We test the hypothesis that PfRH5 opens to insert into the membrane, but instead show that a rigid, disulphide-locked PfRH5 can mediate efficient erythrocyte invasion. Finally, we show that the elongated tail of PfRIPR, which is the target of growth-neutralising antibodies, binds to the PfCSS-PfPTRAMP complex on the parasite membrane. Therefore, a modular PfRIPR is linked to the merozoite membrane through an elongated tail, while its structured core presents PfCyRPA and PfRH5 to interact with erythrocyte receptors. This provides novel insight into the mechanism of erythrocyte invasion and opens the way to new approaches in rational vaccine design.

---

Erythrocyte invasion by *Plasmodium falciparum* involves a tightly ordered sequence of events, starting when the merozoite form of the parasite contacts an erythrocyte^1, 2^. This is followed by a strong actin-dependent deformation of the erythrocyte surface and by reorientation of the merozoite to place its apical pole adjacent to the erythrocyte membrane. Discharge of apical organelles releases the machinery required for invasion, including the PfRCR complex. This leads to a calcium spike at the merozoite-erythrocyte contact site and to the formation of a moving junction between the merozoite and erythrocyte. The parasite then actively pulls its way inside the erythrocyte, followed immediately by a series of deformations of the infected erythrocyte and the establishment of the parasite inside a vacuole.

These stages of erythrocyte invasion require multiple host-parasite interactions, many of which are mediated by protein families with redundant function^1^. However, no strain of *Plasmodium falciparum* has yet been found which can invade erythrocytes when the interaction between PfRH5^3^ and membrane complexes containing the erythrocyte receptor basigin^3–5^ is prevented. Indeed, both PfRH5 and basigin are targets of invasion-neutralising antibodies^3, 6–8^, and immunisation with PfRH5 is protective in an *Aotus* model of malaria^9^ and delays the onset of symptoms in a human challenge model^10^. PfRH5 assembles with PfCyRPA and PfRIPR to form the tripartite PfRCR complex^11–13^. Both PfCyRPA and PfRIPR are also essential for erythrocyte invasion and are targets of invasion-blocking antibodies^13–18^. More recently, a complex containing two merozoite surface proteins, PfCSS and PfPTRAMP, has been shown to bind to PfRCR^19^. This interaction between RIPR, CSS and PTRAMP was first identified in *Plasmodium knowlesi*, where these proteins are also essential for invasion^20^. Invasion-neutralising nanobodies have been identified against both PfCSS and PfPTRAMP^19^ and all five members of PfPCRCR are potential components for a blood-stage malaria vaccine.

Despite the essential function of each component of the PfRCR complex, their roles during invasion are enigmatic. Blocking the function of any member of PfPCRCR halts invasion of erythrocytes at the same step within the invasion process, preventing the increase in calcium concentration at the parasite-erythrocyte interface, an event thought to occur prior to moving junction formation^19^. Several models for PfRH5 function have been proposed, including that PfRH5 and PfRIPR undergo substantial conformational changes to insert into the erythrocyte membrane, forming pores^21^, or that PfRH5 modulates signalling pathways leading to cytoskeletal changes during invasion^22^. However, neither of these hypotheses are robustly supported. To further understand the role of PfRCR in invasion, we therefore determined its structure to 2.89 Å resolution using cryogenic-electron microscopy (cryo-EM) and assessed the role of PfRH5 and PfRIPR in invasion.

## Determining the structure of the PfRCR complex

While crystal structures are available for both PfRH5^5, 6^ and PfCyRPA^14^, structural information for PfRCR was limited to a cryo-EM map of the complex at ∼7 Å resolution^21^. Higher resolution was not achieved in this prior study as the PfRCR complex adopted preferred orientations when vitrified, preventing uniform imaging, and reducing the resolution of the subsequent three-dimensional reconstruction. Crystal structures of PfRH5 and PfCyRPA could be docked into this map, providing insight into the architecture of PfRCR, but this yielded no atomic-level information into the structure of PfRIPR or the interactions between the PfRCR components. To provide an atomic resolution model of PfRCR, we assessed the ability of Fab fragments from monoclonal antibodies which bind to PfRH5 or PfCyRPA to prevent PfRCR from adopting preferred orientations. A complex of PfRCR bound to the Fab fragment of the invasion-neutralising anti-PfCyRPA antibody Cy.003^14^ (Extended Data Fig. 1a-b) had a sufficiently uniform particle distribution to generate a consensus cryo-EM map with a global resolution of 2.89 Å (Fig. 1a, Extended Data Fig. 1c-d, Extended Data Fig. 2, Extended Data Table 1).

**Figure 1:**
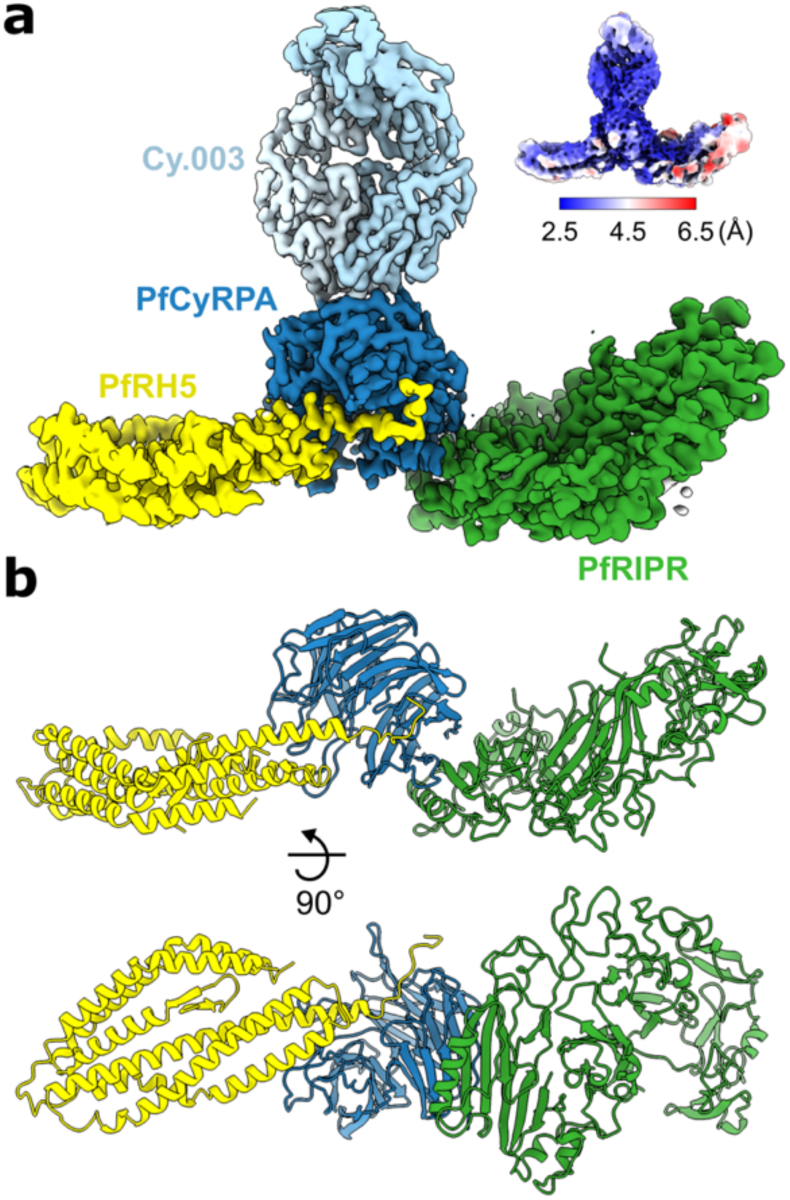
Structure of the PfRCR complex. **a**, Composite map of the PfRCR-Cy.003 complex, following local refinement of the consensus map and post-processing with DeepEMhancer. Density corresponding to PfRH5 (yellow), PfCyRPA (dark blue), PfRIPR (green) and Cy.003 (light blue) are highlighted. The insert is a consensus map prior to local refinement coloured by local resolution. **b**, Structure of the PfRCR complex in cartoon representation, coloured as in **a**. Cy.003 is omitted for clarity.

Crystal structures of PfRH5^5^ and the PfCyRPA-Cy.003 Fab complex^14^ were docked into this map, confirming that PfCyRPA lies at the centre of PfRCR, bridging PfRH5 and PfRIPR (Fig. 1). This arrangement was also consistent with chemical crosslinks observed in crosslinking mass spectrometry (XL-MS) analysis of the PfRCR complex (Extended Data Fig. 3 and Extended Data Table 2). Density attributed to PfRIPR in this consensus map was not large enough to accommodate the full-length molecule, suggesting that part of PfRIPR is flexibly attached to the ordered part of PfRCR and was therefore not resolved. Local resolution estimation indicated that PfCyRPA-Cy.003 was best resolved in the consensus map and PfRIPR the least, especially at its PfCyRPA-distal edge (Fig. 1a).

To investigate the cause of the lower local resolution of PfRIPR, and to identify ways to improve map quality, we subjected particles of the PfRCR-Cy.003 complex to 3D variability analysis in cryoSPARC^23^, revealing conformational heterogeneity (Supplementary Video 1). Flexibility was observed along the length of PfRCR, with both PfRH5 and PfRIPR rocking relative to PfCyRPA and with further internal motion observed within PfRIPR, both consistent with observed local resolution estimates (Fig. 1a). We therefore performed local refinement of both PfRH5 and PfRIPR, significantly improving the density for PfRIPR (Extended Data Fig. 1e and 2). A composite map generated by combining these global and local refinements was used to build a model of PfRCR (Fig. 1b, Extended Data Table 1). Chemical crosslinks identified through XL-MS analysis of the PfRCR complex are consistent with this model (Extended Data Fig. 3 and Extended Data Table 2).

Despite isolating an intact PfRCR-Cy.003 complex by gel filtration (Extended Data Fig. 1a-b), *ab initio* reconstructions showed that a second complex lacking PfRH5 was also present in the dataset (Extended Data Fig. 2). This was separately refined yielding a consensus map with a global resolution of 3.02 Å (Extended Data Fig. 1f-h). The map of this PfCyRPA-PfRIPR-Cy.003 complex also showed lower local resolution at the region corresponding to PfRIPR (Extended Data Fig. 1f), again consistent with flexibility observed in 3D variability analysis. We therefore built a model of PfCyRPA-PfRIPR-Cy.003 complex using a composite map from consensus and local refinements (Extended Data Fig. 4a, Extended Data Table 3).

## Conformational heterogeneity in the PfRH5-PfCyRPA interface

To understand the interface between PfCyRPA and PfRH5, we first assessed the structure of PfRCR-Cy.003, comparing with crystal structures of PfRH5 and PfCyRPA, either free or in complex with ligands or antibodies (Fig. 2a-b and Extended Data Fig. 4). In this interaction, PfCyRPA and PfRH5 each have a buried surface area of ∼1720 Å^2^, with binding mediated by salt-bridges, hydrogen bonds and hydrophobic interactions (Extended Data Table 4). No global conformational changes were observed in PfRH5 or PfCyRPA when compared with the unbound proteins, with much of the PfRH5-PfCyRPA interface formed through rigid body docking of blades 3 and 4 of the PfCyRPA *β*-propeller against helices α5 and α7 of PfRH5 (Fig. 2a). PfRH5 helix α7 projects towards the centre of the β-propeller of PfCyRPA and the most substantial change observed in PfRH5 on PfCyRPA binding is an ordering of its C-terminal tail (residues 507-516). These residues follow α7 and were disordered in previous structures. In PfRCR, they adopt an elongated conformation and occupy a groove which passes between blades 1 and 6 of PfCyRPA, pointing towards but not contacting PfRIPR.

**Figure 2:**
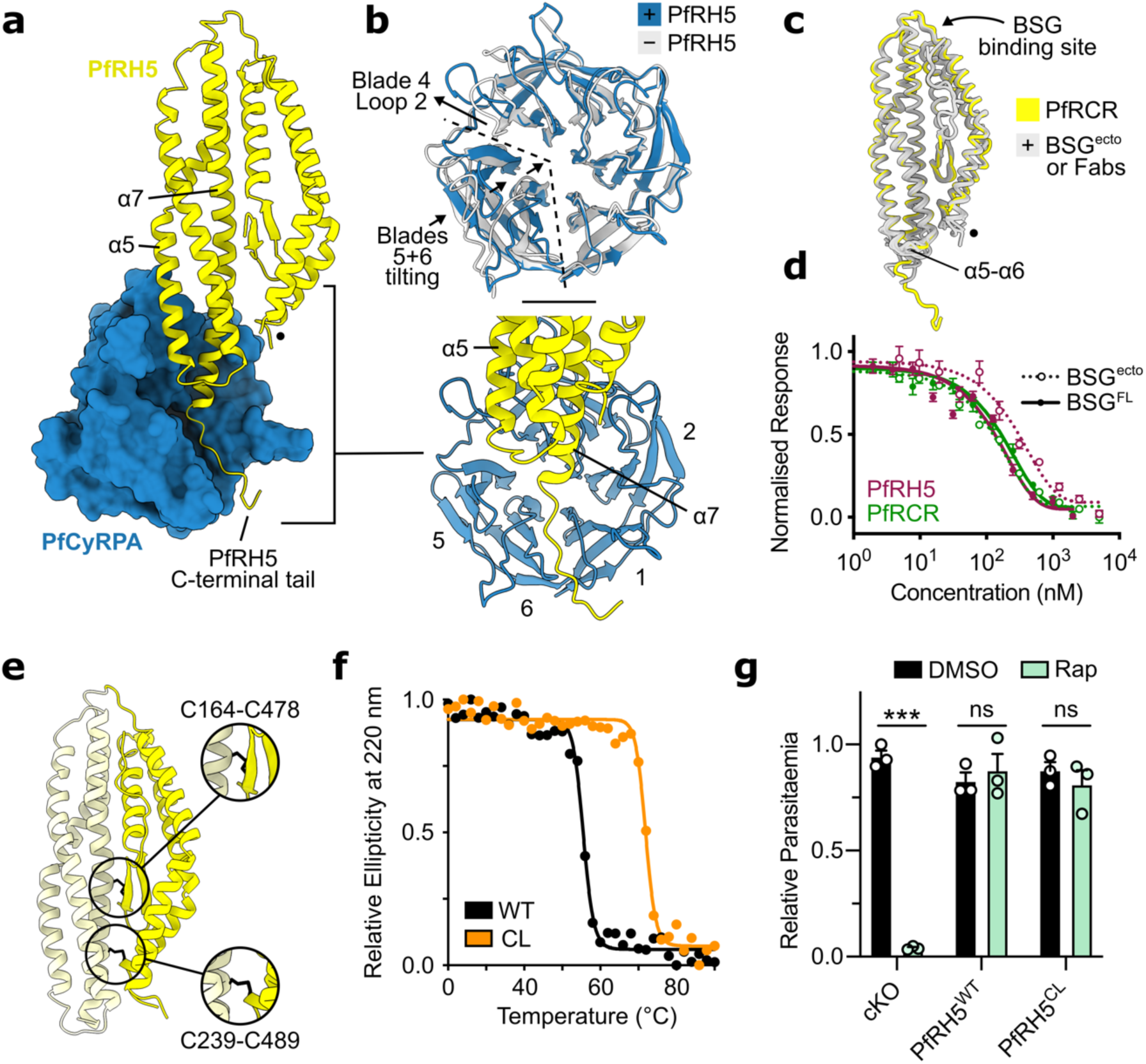
PfRH5 is a rigid component of PfRCR that does not open during invasion. **a**, A view of the interface between PfRH5 (shown as a yellow cartoon) and PfCyRPA (shown as a blue surface). PfCyRPA blade 1 corresponds to residues Ile49-Lys95. The location of the internal loop of PfRH5 between helices α2 and α3, not resolved in this cryo-EM data, is indicated by a black dot. Inset lower right, the interaction between PfRH5 and PfCyRPA is shown in cartoon from the top view with the blades of PfCyRPA labelled. **b**, Overlay of PfCyRPA from structures of PfRCR-Cy.003 (blue) and Cy.003-bound PfCyRPA (grey, PDB ID: 7PI2)^14^, showing structural changes induced by PfRH5 binding. **c**, Overlay of PfRH5 from the PfRCR-Cy.003 structure (yellow) and crystal structures of PfRH5 bound to basigin ectodomain (BSG^ecto^)^5^ or anti-PfRH5 Fab fragments (9AD4, QA1, R5.004, R5.011, R5.015 and R5.016)^5, 6, 14^ (grey). The binding site of basigin, the loop linking α5 and α6, and the location of the internal loop of PfRH5 (black dot) are highlighted. PDB codes used: 4U0Q, 4U0R, 4U1G, 6RCU, 6RCV and 7PHU. **d**, Normalised microscale thermophoresis responses showing equivalent binding of both basigin ectodomain (BSG^ecto^, dotted lines) and full-length basigin (BSG^FL^, full lines) to PfRH5 (maroon) and PfRCR (green). Each sample was measured in triplicate for two independent technical replicates, one of which is shown. The mean and standard deviation are shown. **e**, A model of the designed double cysteine-locked PfRH5 construct (PfRH5^CL^) and its disulphide bonds at Cys164-Cys478 and Cys239-Cys489 (black). These two disulphide bonds lock the N-terminal (dark yellow) and C-terminal (light yellow) halves of PfRH5 together, preventing opening of PfRH5. **f**, The relative ellipticity of wild-type PfRH5 (WT, black) and double cysteine-locked PfRH5 (CL, orange) as a function of temperature, measured by circular dichroism, demonstrating the improved thermal stability of PfRH5^CL^ by ∼14 °C. **g**, Relative parasitaemia of transgenic *P. falciparum* parasites with PfRH5 conditionally knocked out (PfRH5^cKO^) or replaced with wild-type PfRH5 (PfRH5^WT^) or double cysteine-locked PfRH5 (PfRH5^CL^) upon rapamycin (Rap) treatment. Rapamycin treatment of PfRH5^cKO^ causes a significant reduction in parasitaemia (*** indicates p < 0.001) whereas no significant differences in parasitaemia are observed when cKO is complemented with either PfRH5^WT^ or PfRH5^CL^ (p = 0.599 and 0.457, respectively, determined by two-tailed, unpaired t-test). In all cases, transgenic parasite lines show no significant differences in growth in the presence of DMSO only. Parasite growth was measured in biological triplicate, with each of these also measured in technical triplicate. Individual data points, the mean and standard error of the mean are shown.

A more substantial conformational change is observed in PfCyRPA when present in the PfRCR complex (Fig. 2b). While blades 1 to 4 overlay closely with those in crystal structures of PfCyRPA (RMSD 0.633 Å to Cy.003-bound PfCyRPA and 0.662 Å to unbound PfCyRPA (Fig. 2b and Extended Data Fig. 4b)), blades 5 and 6 and the linker between these blades move closer to the central axis of its β-propeller when in PfRCR, predominantly through a rigid-body tilt. This alters the width of the groove within which residues 507-516 of PfRH5 lie. This is accompanied by a substantial and more localised rearrangement in loop 2 of blade 4, which moves by 11.5 Å at Val250 to accommodate helix α5 of PfRH5, with additional changes to loop 2 of each of blades 1 to 3 and to the linker between blades 3 and 4. To determine whether these changes were induced by PfRH5 binding, we compared the structure of PfCyRPA in PfRCR-Cy.003 with that in the PfCyRPA-PfRIPR-Cy.003 complex (Extended Data Fig. 4b). In this latter complex, PfCyRPA adopts a structure more like that observed in crystal structures (with RMSDs of 0.629 Å to Cy.003-bound PfCyRPA and 0.622 Å to unbound PfCyRPA, versus 0.858 Å to PfCyRPA in PfRCR-Cy.003), indicating that these conformational changes are the result of PfRH5 binding. Notably, in both PfRH5 and PfCyRPA, the epitopes previously found to be the targets of neutralising antibodies^5, 6, 14^ do not change in conformation after complex formation, supporting vaccine immunogen design approaches based on original crystal structures.

## PfRH5 forms a rigid structure which does not open during erythrocyte invasion

Previous studies have suggested that PfRH5 is flexible, that its binding to basigin is influenced by its assembly into the PfRCR complex, and that the N- and C-terminal halves of PfRH5 open to allow it to insert into the erythrocyte membrane to form a pore^21^. To test these hypotheses, we analysed the structure of PfRH5 from the PfRCR-Cy.003 complex. Due to global rigid body motions along the length of the PfRCR complex (Supplementary Video 1), the PfCyRPA-distal tip of PfRH5 is one of the lower resolution regions of the cryo-EM map, as previously observed^21^, and as expected for the extremities of a flexible protein complex structurally characterised through single particle averaging. However, after local refinement to partially correct this flexibility, we found that the PfCyRPA-distal tip of PfRH5 had a local resolution of ∼4-5 Å, sufficient for a molecular model to be reliably built (Extended Data Fig. 4c-d). Comparison of this model with crystal structures of PfRH5, either bound to basigin^5^, or to monoclonal Fab fragments^5, 6, 14^ revealed no consistent conformational changes in PfRH5 as a result of being part of PfRCR (with RMSDs to each of less than 1 Å) (Fig. 2c, Extended Data Fig. 4e). The only variable regions of PfRH5 across this alignment are the loop linking helices α5 and 6 and the C-terminal end of α7. These are both within the region of PfRH5 which contacts PfCyRPA. The PfCyRPA-bound conformation is not an outlier when compared with other PfRH5 structures, suggesting that these are intrinsically flexible regions of PfRH5 which become ordered upon binding to PfCyRPA. We therefore observe no conformational change in PfRH5 induced by binding to PfCyRPA.

The finding that PfRH5 adopts a similar conformation in PfRCR as when bound to basigin^5^, led us to question the observation that PfRCR does not bind to basigin ectodomain^21^. We therefore measured binding of PfRCR and PfRH5 to either basigin ectodomain (BSG^ecto^) or to detergent solubilised full-length basigin (BSG^FL^) using microscale thermophoresis (Fig. 2d, Extended Data Fig. 4f). Binding was observed in each case, with affinities ranging from ∼0.12 μM for PfRH5 binding to full-length basigin to ∼0.19 μM for PfRH5 to basigin ectodomain, with the affinities for PfRCR binding to either form of basigin falling between these values (Extended Data Fig. 4f). We therefore find that PfRCR and PfRH5 bind to basigin and to basigin ectodomain with similar affinities, not supporting the hypothesis that assembly into the PfRCR complex changes the basigin binding properties of PfRH5.

We next tested the hypothesis that the N- and C-terminal halves of PfRH5 might come apart during the erythrocyte invasion process, allowing PfRH5 to form a pore which causes a spike in calcium concentration at the merozoite-erythrocyte interface^21^. To achieve this, we introduced locking disulphide bonds into the centre of PfRH5 which would prevent such a conformational change and used a conditional replacement system to introduce these into a *P. falciparum* line to assess their effect on erythrocyte invasion (Fig. 2e-g). We employed a Rosetta^24^-based, structure-guided approach to design five disulphide bonds which we predicted would hold together the N- and C-terminal halves of PfRH5 (Cys164-Cys478 (CC1), Cys180-Cys471 (CC2), Cys300-Cys408 (CC3), Cys166-Cys481 (CC4) and Cys239-Cys489 (CC5)) (Extended Data Fig. 5a). In each case, these were introduced into a version of PfRH5 lacking the N-terminus and disordered loop (PfRH5ΔNL)^5^, were expressed in *Drosophila* S2 cells and were shown by circular dichroism analysis to adopt the expected secondary structure (Extended Data Fig. 5b-d). We used circular dichroism with a thermal melt to test the stability of these five variants and found that two of the designed disulphide bonds increased the thermal stability of PfRH5ΔNL by greater than 5 °C, as would be expected on successful formation of a stabilising disulphide bond (+6 °C for Cys164-Cys478 and +8 °C for Cys239- Cys489) (Extended Data Fig. 5e). These two disulphide bonds were then combined to generate a final cysteine-locked PfRH5 design (PfRH5^CL^) (Fig. 2e, Extended Data Fig. 5c and f), and this combination increased the thermal stability of PfRH5ΔNL by ∼14 °C, indicating that both designed cysteine locks were formed (Fig. 2f). We further validated the presence of these disulphide bonds in PfRH5^CL^ by mass spectrometry analysis following maleimide-PEG2-biotin labelling^25^, which modifies free sulfhydryl groups, but not disulphide-bonded cysteine residues. Non-reduced PfRH5ΔNL^CL^ was modified by the addition of just one maleimide, indicating that it contains only one free cysteine, most likely corresponding to the native single free cysteine, Cys239 (Extended Data Fig. 5g). In contrast, reduced PfRH5ΔNL^CL^ showed a series of species following labelling, containing from zero to eight maleimide modifications, consistent with a total of nine cysteine residues and partial labelling. Together these data confirm that PfRH5^CL^ contains four disulphide bonds (two native and two designed) which are formed under non-reducing conditions.

We next generated a transgenic *P. falciparum* line which could be used to induce a conditional replacement of the native PfRH5 gene with the cysteine-locked variant (Extended Data Fig. 6). To achieve this, *LoxP* sites were inserted silently within a synthetic intron^26^ either side of exon 2 of the endogenous PfRH5 gene, generating a conditional knockout line (PfRH5^cKO^) in which the addition of rapamycin would lead to excision of the coding sequence of the gene (Extended Data Fig. 6a). Two more strains were then generated in which a second copy of PfRH5 (either a wild-type sequence, PfRH5^WT^ or the cysteine-locked variant, PfRH5^CL^) was inserted downstream of the endogenous floxed PfRH5 gene, such that addition of rapamycin would lead to removal of the endogenous PfRH5 sequence and expression of the downstream copy (Extended Data Fig. 6b-e). We then tested the ability of all three lines to invade erythrocytes by treating ring stage parasites with rapamycin or DMSO as a control, then measuring their resultant parasitaemia (Fig. 2g). The addition of rapamycin to PfRH5^cKO^, which lacks a downstream replacement copy of PfRH5, led to a >95 % reduction in parasitaemia, confirming the requirement of PfRH5 for invasion. However, in parasite lines with a downstream copy of either native PfRH5^WT^ or PfRH5^CL^, the addition of rapamycin did not affect parasitaemia, demonstrating that both were able to complement the deletion of PfRH5 and allow effective erythrocyte invasion. The ability of a PfRH5 variant which is locked through two disulphide bonds across its N- and C-halves to mediate invasion is not compatible with a model in which the PfRH5 structure is required to ‘open’ such that it can insert into the erythrocyte membrane. Instead, this is consistent with PfRH5 adopting a rigid structure throughout the process of erythrocyte invasion.

## The structure of PfRIPR

We next used our composite cryo-EM map of PfRCR-Cy.003 to build a molecular model of PfRIPR (Fig. 3). The map resolution was sufficient to build much of the structure *de novo* (Extended Data Fig. 7a), with molecular models derived from structure prediction using a local installation of AlphaFold2^27^ used to guide building (Extended Data Fig 7b), particularly in the less well-ordered regions of the map. This resulted in an almost complete model of residues 34-716 of PfRIPR (which has a total length of 1086 residues), with no density observed for loops comprising residues 124-137 and 479-557 (Fig. 3). Moreover, we did not observe interpretable density for residues 717-1086 of PfRIPR, revealing that this C-terminal region of PfRIPR is flexibly connected to the PfRCR complex. This was consistent with the internal crosslinks found within PfRIPR from XL-MS analysis of PfRCR, most of which were in the first ∼700 residues (Extended Data Fig. 3).

**Figure 3:**
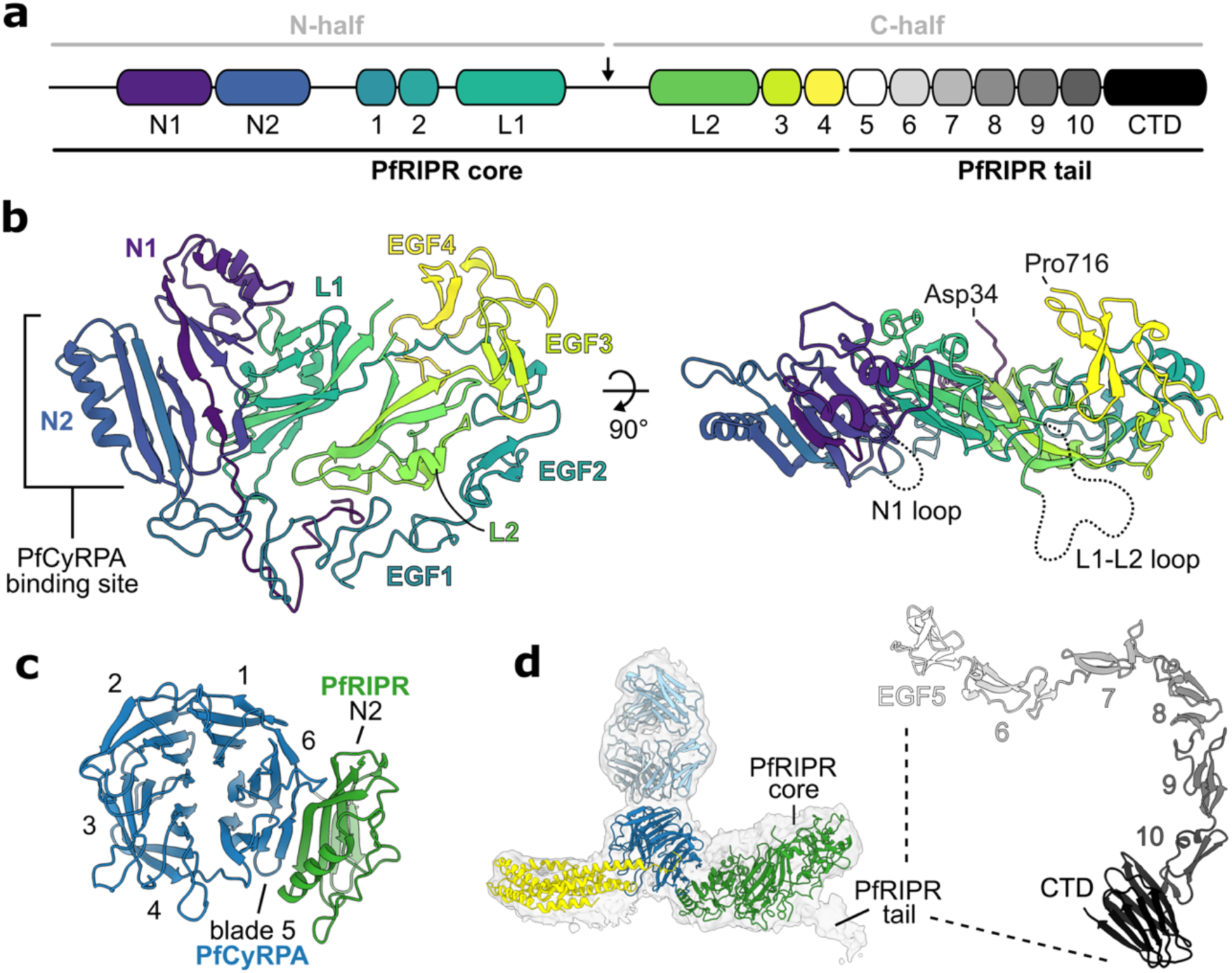
The structure of PfRIPR. **a**, The domain architecture of PfRIPR. The PfRIPR core comprises the N-terminus to EGF-like domain 4, and the PfRIPR tail comprises EGF-like domain 5 to the C-terminus. In addition to the previously known ten EGF-like domains, PfRIPR contains two N-terminal domains N1 and N2, two lectin-like domains L1 and L2, and a C-terminal domain (CTD). PfRIPR is cleaved at a site in the L1-L2 loop to produce its N- and C-halves, indicated by an arrow. **b**, Structure of the PfRIPR core (residues Asp34-Pro716) coloured sequentially from its N-terminus (blue) to C-terminus (yellow), as in panel **a**. The location of the PfCyRPA binding site on domain N2, and the unresolved N1 loop (residues 124-137) and L1-L2 loop (residues 479-557) are shown. **c**, A view of the interface between PfCyRPA and PfRIPR, predominantly composed of an extended beta-sheet between PfCyRPA blade 5 and PfRIPR domain N2, plus some interactions with the helix of N2. Only domain N2 of PfRIPR is shown for clarity. **d**, Map of the PfRCR-Cy.003 complex containing additional density for the beginning of the PfRIPR tail. The AlphaFold2 prediction of the PfRIPR tail is shown (right) with the EGF-like domains and the CTD labelled and coloured in greyscale as in **a**.

The structure of PfRIPR allowed us to re-define its domain architecture (Fig. 3a,b). PfRIPR has previously been described as consisting of two halves generated by cleavage of PfRIPR^28^, with these two halves remaining associated^13^. Through sequence analysis, PfRIPR was also predicted to contain ten EGF-like domains, with two in the N-half (EGF1 and EGF2) and eight in the C-half (EGF3-10)^13^. In our PfRIPR structure, the predicted cleavage site that joins the two predicted halves of PfRIPR is found within a large unstructured loop (residues 479-557). However, this loop does not link two separate halves of PfRIPR, but instead emerges from within an ordered array of different domains, which ends after domain EGF4. We call this region the PfRIPR core.

The core of PfRIPR adopts a complex and novel architecture in which eight domains are intertwined (Fig. 3b). The two N-terminal domains, N1 and N2, each consist of a three or four- stranded *β*-sheet packing against one or two α-helices. These two domains interact with one another through their *β*-sheets and it is through domain N2 that PfRIPR binds to PfCyRPA, with the three stranded *β*-sheet of N2 forming a continuation of the *β*-sheet of PfCyRPA blade 5 (Fig. 3c). Some additional contacts are formed between the long *α*-helix of N2 and the face of the same PfCyRPA blade, and between domain N1 and the N-terminal *β*-strand of PfCyRPA, which is found in blade 6 (Extended Data Table 4). After a short sequence following N2, the first two EGF-like domains of PfRIPR, EGF1 and EGF2 follow, wrapping around one edge of the core. Next are two domains, L1 and L2, which form the centre of the PfRIPR core, and around which the other domains fold. Intriguingly, L1 and L2 adopt the same fold as rhamnose-binding lectins and therefore the same fold as PfP113^29^ (Extended Data Fig. 7c), a proposed binding partner of PfRH5 that was previously thought to anchor PfRH5 to the merozoite surface^30^, but which was recently disputed as a surface protein^31^. The large disordered internal loop where PfRIPR is cleaved links L1 and L2. Multiple interdomain contacts within the PfRIPR core, including those between L1 and L2, hold the N-half and C-half of PfRIPR together following loop cleavage. Finally, EGF3 and EGF4 pack against the linker which joins EGF2 and L1, wrapping around the opposite edge of the PfRIPR core. This leaves the C-terminus of the PfRIPR core at the side opposite to the PfCyRPA binding site, and on the opposing face to the N1 and L1-L2 loop. While the PfRIPR core is cysteine-rich, containing twenty-three disulphide bonds and one free cysteine (Cys256), these are all contained within domains and there are no interdomain disulphide bonds (Extended Data Fig. 7d).

While the map of PfRCR-Cy.003 did not contain density of sufficient resolution to allow model building of PfRIPR beyond residue 716, in several 2D classes we could see weaker additional density projecting away from the PfRIPR core (Extended Data Fig. 2). This density could not be improved or extended using 2D classification with a larger particle box size, suggesting that it results from residues flexibly linked to the PfRIPR core. Nonetheless, non-uniform refinement of a subset of particles yielded a map at a global resolution of 3.95 Å (Fig. 3d and Extended Data Fig. 7e), which showed that this extra density projects away from the centre of the PfRIPR core, adjacent to where EGF4 ends at residue 716. This suggested that the rest of PfRIPR extends away from the PfRIPR core at this location. Indeed, XL-MS analysis identified a crosslink between Lys736 of EGF5 and Lys427 of domain L1, indicating that EGF5 is located approximately in this area (Extended Data Fig. 3 and Extended Data Table 2), and as supported by an AlphaFold2 prediction of PfRIPR up to the end of EGF5 (Extended Data Fig. 7f). To generate a complete composite model of full-length PfRIPR, we used AlphaFold2 to predict the structure of residues 717 to 1086 of PfRIPR. This predicts an elongated but ordered structure, which we call the PfRIPR tail, consisting of EGF-like domains 5-10 followed by a C-terminal domain (CTD) with a Galectin-like fold (Fig. 3d, Extended Data Fig 7c). Guided by the additional density in the 3.95 Å map, we docked this model of the PfRIPR tail onto the PfRIPR core to generate a complete molecular model for PfRIPR (Extended Data Fig. 7g).

## The PfRIPR tail binds the PfCSS-PfPTRAMP complex allowing PfRIPR to bridge the merozoite and erythrocyte membranes

The *Plasmodium knowlesi* homologue of PfRIPR, PkRIPR, and more recently PfRIPR itself, have been shown to bind their respective homologues of the CSS-PTRAMP complex^19, 20^. We therefore asked whether it is the core or the tail of PfRIPR that mediates binding to PfCSS- PfPTRAMP. We immobilised full-length PfRIPR and the PfRIPR tail (Extended Data Fig. 8a) on two flow paths of a surface plasmon resonance chip, then flowed the PfCSS-PfPTRAMP complex over these surfaces (Extended Data Fig. 8b). Both PfRIPR and PfRIPR tail bound to PfCSS-PfPTRAMP with similar affinities of ∼3 μM (Fig. 4a, Extended Data Fig. 8c-d), indicating that the PfRIPR tail alone is sufficient for binding to the transmembrane-anchored PfCSS- PfPTRAMP complex. This positions the complex containing the PfRIPR core, PfCyRPA and PfRH5 such that it can bind to receptors, like basigin, on the erythrocyte surface.

**Figure 4:**
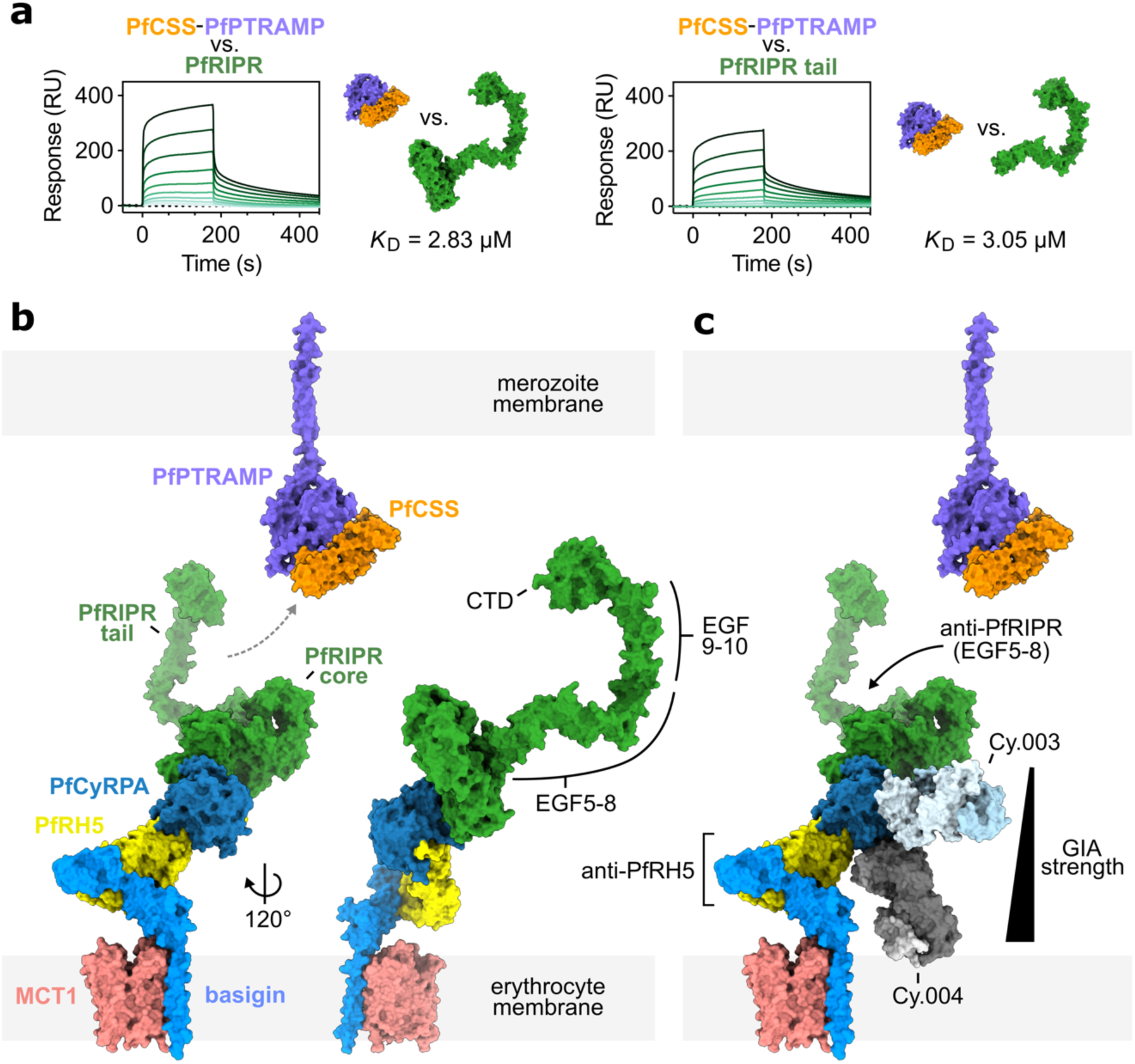
The PfPCRCR complex bridges the parasite and erythrocyte membranes. **a**, Surface plasmon resonance curves showing that the PfCSS-PfPTRAMP complex binds to both full-length PfRIPR and the PfRIPR tail with equal affinity. Curves show binding of a two- fold dilution series of the PfCSS-PfPTRAMP complex, representative of data measured in triplicate. The binding affinity is the mean of those determined by steady state analysis of each independent measurement (Extended Data Figure 8c-d). Models of PfCSS, PfPTRAMP ectodomain, composite full-length PfRIPR and PfRIPR tail are shown inset, as described in (**b**). **b**, Composite model of the PfRCR complex on the erythrocyte membrane, illustrating that the tail of PfRIPR projects towards the merozoite membrane where the apical membrane-bound PfCSS-PfPTRAMP complex is located. EGF-like domains 5-8, 9-10 and the C-terminal domain (CTD) of PfRIPR are highlighted. The composite PfRCR complex model is aligned on to the structure of basigin-bound MCT1 (PDB ID: 7CKR^32^). No structure of the PfCSS-PfPTRAMP complex has been solved; for illustrative purposes only, the AlphaFold2 predicted structure of PfPTRAMP (magenta, AlphaFoldDB entry: Q8I5M8, residues 26-352) and the crystal structure of PfCSS (orange, PDB ID: 7UNY^19^) are therefore docked upon one another. **c**, Mapping of the growth-inhibitory monoclonal antibody Fab fragments targeting PfRCR components on the complex in context of binding to the erythrocyte membrane. Growth- inhibitory PfRH5-targeting antibodies sterically interfere with the binding of PfRH5 to membrane-bound basigin. The strength of growth inhibitory PfCyRPA-targeting antibodies correlates with their proximity to the erythrocyte membrane, illustrated by overlay of the Cy.004 Fab-bound PfCyRPA crystal structure (PDB ID: 7PHW^14^) on to the PfRCR-Cy.003 complex. The epitopes of growth-inhibitory PfRIPR-targeting antibodies are found in EGF-like domains 5-8, located on the opposite surface of the PfRCR complex, distal to the erythrocyte membrane.

## Understanding the mechanisms of action of invasion-neutralising antibodies

Invasion-neutralising monoclonal antibodies have been identified against each of PfRH5, PfCyRPA and PfRIPR. To understand how these antibodies block the invasion process, we mapped their binding sites onto a model of PfRCR. As erythrocyte basigin is all found in a stable complex with either plasma membrane calcium transporters (PMCAs) or with the monocarboxylate transporter (MCT1)^4^, we combined our structure of PfRCR with that of the PfRH5-basigin complex^5^ and with structures of basigin-MCT1^32^ and basigin-PMCA^33^ to generate composite models of the PfRCR complex on the erythrocyte surface (Fig. 4b and Extended Data Fig. 8e). While there will be some flexibility in the presentation of the PfRCR complex due to hinge movements in basigin, modelling predicts that PfRCR will project away from the erythrocyte surface, with PfRH5 closest to the erythrocyte and with the PfRIPR tail projecting towards the merozoite surface. This arrangement is consistent with surface plasmon resonance binding data (Fig. 4a), which indicates that the PfRIPR tail mediates binding to PfCSS-PfPTRAMP.

PfRH5- and PfCyRPA-targeting neutralising antibodies are proposed to act by sterically blocking the approach of PfRCR to the erythrocyte membrane, preventing its binding to basigin and any potentially unknown surface receptors^14, 18^. Indeed, the most effective PfRH5- binding neutralising antibodies block PfRH5 from binding to basigin-PMCA and basigin-MCT1 complexes^4^. There is no known erythrocyte binding partner for PfCyRPA and the mechanism by which invasion blocking anti-PfCyRPA antibodies act is uncertain. Here, we find that when aligned onto the composite model of PfRCR-basigin-MCT1 and PfRCR-basigin-PMCA, the degree to which these antibodies project towards the erythrocyte membrane correlates with their growth-inhibitory activity^14^ (Fig. 4c and Extended Data Fig. 8e-f). Aligning the crystal structure of PfCyRPA bound to the strongly neutralising antibody Cy.004^14^ onto PfRCR places the Fab fragment of Cy.004 projecting towards the erythrocyte surface, where it would sterically clash with the membrane when part of an intact monoclonal antibody. In contrast, the less neutralising antibodies Cy.003, Cy.007 and c12 lie almost parallel to the erythrocyte membrane, and 8A7 lies between Cy.003 and Cy.004. The invasion-blocking activity of these antibodies^14^ thus correlates with the degree to which they project towards the erythrocyte membrane.

There are fewer studies of growth-inhibitory antibodies that target PfRIPR. As PfRCR projects away from the erythrocyte surface, antibodies which bind to the PfRIPR core are unlikely to prevent binding to erythrocytes. Indeed, the most effective known polyclonal and monoclonal antibodies targeting PfRIPR bind to EGF-like domains 5 to 8 in the PfRIPR tail^13, 17, 18^. How these antibodies block the invasion process will require further study but, as the PfRIPR tail binds to PfCSS-PfPTRAMP on the merozoite surface, it is possible that they affect processes such as receptor engagement on the merozoite surface rather than on the erythrocyte surface.

## Conclusions

Our cryo-EM structure of PfRCR has allowed us to build a first atomic model of PfRIPR, revealing it to consist of a complex multi-domain core flexibly attached to an elongated tail, containing domains which adopt the same fold as galectin- and rhamnose-binding lectins. We reveal the conformations of PfRH5, PfCyRPA and PfRIPR when part of PfRCR, and define their interactions at a molecular level. We find that PfRH5 does not differ substantially in structure when integrated into the PfRCR complex, and that disulphide-locking the two halves of PfRH5 together has no impact on erythrocyte invasion, indicating that PfRH5 does not open to insert into the erythrocyte membrane as part of the invasion process. Additionally, we find that the tail of PfRIPR binds to the PfCSS-PfPTRAMP complex, which is anchored at the merozoite surface, suggesting that PfRCR bridges the erythrocyte and parasite membranes during invasion. The mapping of growth-neutralising antibodies on to the structure of erythrocyte-bound PfRCR supports a model in which antibodies which target PfRH5 and PfCyRPA sterically prevent PfRCR from binding to the erythrocyte membrane, while the most effective PfRIPR- targeting invasion-neutralising antibodies may act towards the parasite membrane. These studies inform and open new future avenues to understand the molecular mechanisms of erythrocyte invasion and the rational design of blood-stage malaria vaccines.

## Methods

### Protein expression and purification

PfRH5 (residues E26-Q256, with substitutions C203Y (of the 7G8 *P. falciparum* strain) and T216A and T299A (to remove potential N-linked glycosylation sites), a BiP secretion sequence and a C-terminal C-tag) was expressed from a stable S2 cell line^34^ (ExpreS^2^ion Biotechnologies) in EX-CELL® 420 Serum Free Medium (Sigma Aldrich). After 3-4 days, culture supernatants were harvested and 0.45 µm filtered, then incubated with CaptureSelect™ C-tagXL resin (Thermo Scientific). Beads were washed using 20-30 column volumes of TBS (25 mM Tris pH 7.5, 150 mM NaCl) and bound proteins eluted with C-tag elution buffer (25 mM Tris pH 7.5, 2 M MgCl2). Eluted proteins were further purified by gel filtration using a S200 Increase 10/300 column with a running buffer of HBS (20 mM HEPES pH 7.5, 150 mM NaCl).

PfCyRPA (residues D29-E362, with substitutions S147A, T324A and T340A (to remove potential N-linked glycosylation sites), a mammalian secretion sequence and a C-terminal C- tag) was transiently expressed using Expi293F™ cells using the Expi293™ Expression System Kit (Thermo Fisher) as recommended. Culture supernatants were harvested and 0.45 µm filtered, diluted 1:1 in TBS and then incubated with CaptureSelect™ C-tagXL resin. The purification then proceeded as for PfRH5.

PfRIPR (residues D21-N1086, with substitutions N103Q, N144Q, N228Q, N303Q, N334Q, N480Q, N498Q, N506Q, N526Q, N646Q, N647Q, N964Q and N1021Q (to remove potential N-linked glycosylation sites), a BiP secretion sequence and a C-terminal C-tag) was expressed from a stable S2 cell line as for PfRH5. After 3-4 days, culture supernatants were harvested and 0.45 µm filtered before being concentrated and buffer-exchanged into 50 mM Tris pH 7.5, 150 mM NaCl by tangential flow filtration with a stack of three 100 kDa Omega™ Cassettes (PALL Corporation). The exchanged supernatant was loaded on to a 1 ml pre-packed CaptureSelect™ C-tagXL column (Thermo Scientific) equilibrated with TBS. After washing with 30-50 column volumes of TBS, bound proteins were eluted with C-tag elution buffer. Eluted proteins were exchanged into HBS using a PD-10 desalting column (Cytiva), then either further purified by gel filtration as for PfRH5 or used directly for PfRCR complex preparation.

PfRIPR tail (residues D717-N1086, with substitutions N964Q and N1021Q (to remove potential N-linked glycosylation sites), a BiP secretion sequence and a C-terminal C-tag) was also expressed from a stable S2 cell line. PfRIPR tail was purified as for full-length PfRIPR except that filtered culture supernatants were exchanged into TBS by tangential flow filtration with a stack of three 5 kDa Omega™ Cassettes (PALL Corporation). Eluted proteins were further purified by gel filtration using a S200 Increase 10/300 column into HBS.

PfCSS (residues Q21-K290, with a secretion sequence and C-terminal His6-tag) and PfPTRAMP (residues C30-T307, with a secretion sequence and C-terminal His6-tag) were transiently expressed using FreeStyle™ 293-F cells (Thermo Fisher) in FreeStyle™ F17 Expression Medium supplemented with L-glutamine and 1X MEM non-essential amino acids (Gibco). 6 days following transfection, culture supernatants were harvested and 0.45 µm filtered, then incubated with Super Ni-NTA resin (Generon) equilibrated in HBS. After washing with 20 column volumes of HBS supplemented with 20 mM imidazole, bound proteins were eluted with HBS supplemented with 300 mM imidazole, then further purified by gel filtration using an S75 Increase 10/300 column into HBS.

Cy.003^14, 35^ was transiently expressed in Expi293F™ cells. Culture supernatants were harvested and 0.45 µm filtered before loading on to a 1 ml HiTrap™ Protein G HP column (Cytiva) pre-equilibrated in PBS (79382, Sigma-Aldrich). The column was washed with PBS and bound proteins were eluted using 5 ml of 0.1 M glycine pH 2.5 into 1 ml of 1 M Tris pH 8. To prepare Fab fragments, the eluted mAb was cleaved using Immobilised Papain (20341, Thermo Scientific). After cleavage, Fc and Fab fragments were separated using a 1 ml HiTrap™ rProtein A pre-packed column (Cytiva), and the unbound fraction that contained Cy.003 Fab was exchanged into PBS.

To express full-length human basigin, a synthetic gene (UniProt ID P35613-2) was cloned into pFastBac with a C-terminal His6-tag for expression in Sf9 insect cells using the Bac-to-Bac™ Baculovirus Expression System (Thermo Fisher). Following transformation of DH10Bac cells, bacmids were isolated by isopropanol precipitation and used to transfect Sf9 cells at a cell density of 1 million cells/ml in Sf-900™ II serum-free media (Gibco). First generation baculoviruses (P1) were amplified to produce second generation baculoviruses (P2), then these were used to induce expression of full-length basigin by addition at 1% v/v to Sf9 cells at ∼2.5-3.0 million cells/ml. After 48 hours, cells were harvested then resuspended in lysis buffer (25 mM Tris pH 8.0, 150 mM NaCl, 10% glycerol) supplemented with cOmplete™ EDTA- free protease inhibitors (Roche). Cells were lysed using a Dounce homogeniser followed by sonication on ice (60% amplitude, 3 seconds pulse, 9 seconds rest, for 1.5 mins). Lysed homogenate was clarified by centrifugation at 3,000 g for 20 mins, then the supernatant further spun at 100,000 g for 45 mins at 4 °C to isolate membranes. After resuspension in lysis buffer, membranes were solubilised with 1.2% DDM for 1 hour at 4 °C. After centrifugation at 100,000 g for 45 mins at 4 °C, the supernatant with incubated with Ni-NTA resin (Qiagen) at 4 °C for 1 hour, then the resin washed with 25 mM Tris pH 8.0, 300 mM NaCl, 10% glycerol, 0.02% DDM and 15 mM imidazole. Bound proteins were eluted using 25 mM Tris pH 8.0, 150 mM NaCl, 10% glycerol, 0.02% DDM, 400 mM imidazole, then further purified on an S200 Increase 10/300 column into buffer containing 20 mM HEPES pH 7.2, 150 mM NaCl and 0.02% DDM:CHS (10:1 w/v ratio). Approximately 100-200 µg of full-length basigin was obtained from 100 ml of Sf9 cell culture.

Basigin ectodomain was expressed and purified as reported previously^5^.

### Structure determination using cryogenic electron microscopy

To prepare a complex of PfRH5-PfCyRPA-PfRIPR-Cy.003 Fab for cryo-EM analysis, purified proteins were incubated together at an equimolar ratio in HBS for 5 minutes at room temperature. Approximately 250 µg of complex was prepared. After incubation, the mixture was subject to gel filtration using an S200 Increase 10/300 GL column equilibrated in HBS. Fractions containing the complex were concentrated with a 100K Amicon® Ultra centrifugal unit at 6,000 g and 4 °C.

Cryo-EM grids were prepared with an FEI Vitrobot Mark IV (Thermo Fisher) at 4 °C and 100% humidity. 3 µl of complex at 0.2 mg/ml was applied to Au-Flat 1.2/1.3 grids (Protochips) which had been glow-discharged at 15 mA for 60 seconds. After incubating for 5 seconds, grids were blotted for 1 to 4 seconds then plunged into liquid ethane.

Grids were imaged using an FEI Titan Krios operating at 300 kV and equipped with a Gatan BioQuantum energy filter (20 eV) and K3 direct electron detector. Data collection was automated using the fast acquisition mode in EPU (Thermo Fisher). Images were acquired at a nominal 58,149x magnification corresponding to a calibrated pixel size of 0.832 Å/pixel (0.416 Å/super-resolution pixel), with a dose rate of 16.32 electrons/Å^2^/second, and a total exposure time of 3 seconds with 40 frames. This resulted in a total dose of 48.97 electrons/Å^2^. Images were acquired using a 100 µm objective aperture and a defocus range of -1.0 to -3.0 µm in 0.25 µm increments. Data were collected from 3 grids in back-to-back sessions, all prepared with the same sample. A total of 13,542 movies were acquired (7,428 from grid 1, 2,720 from grid 2 and 3,376 from grid 3).

### Image processing

Movies were motion corrected and CTF parameters were estimated on-the-fly in SIMPLE 3.0^36^. Datasets from each session were pre-processed individually. Micrographs were first template-picked using templates from a previous pilot data collection and picked particles (1,202,046 from grid 1, 1,601,557 from grid 2, and 752,273 from grid 3 – yielding a total of 2,555,876 particles) were extracted (box size of 416 pixels) and subject to 2D classification separately. After excluding particles from poorly defined 2D classes, particles from all three sessions were exported to cryoSPARC v3.3.2. Here, particles were subject to further rounds of 2D classification and particle clean-up, yielding a combined total of 961,077 particles. These were used for *ab initio* reconstruction into 6 classes, which revealed that the dataset contained three major species: the PfRCR-Cy.003 complex, a complex lacking PfRH5 (PfCyRPA-PfRIPR-Cy.003), and a complex of mostly PfCyRPA and Cy.003 alone.

Before further refinement, micrographs were repicked using the cryoSPARC implementation of TOPAZ^37^, yielding a total of 2,615,684 particles. These were subject to rounds of 2D classification to remove bad particles, then combined with the previous SIMPLE-picked particles. After removal of duplicates, a final particle stack of 1,686,994 unique particles was obtained. These particles were subject to heterogeneous refinement using volumes for the PfRCR-Cy.003, PfCyRPA-PfRIPR-Cy.003 and PfCyRPA-Cy.003 complexes, plus three decoy volumes. This separated the particles into PfRCR-Cy.003 (523,352 particles), PfCyRPA-PfRIPR- Cy.003 (527,499 particles), PfCyRPA-Cy.003 (413,374 particles), decoy 1 (107,015 particles), decoy 2 (97,538 particles), and decoy 3 (18,216 particles). Further homogeneous then non-uniform refinement of PfRCR-Cy.003 and PfCyRPA-PfRIPR-Cy.003 particles yielded maps of 3.15 Å for PfRCR-Cy.003 and 3.27 Å for PfCyRPA-PfRIPR-Cy.003. After Bayesian polishing of particles in RELION 3.1.3 using the default settings^38^ and local per-particle CTF refinement in cryoSPARC, a final non-uniform refinement yielded consensus maps of 2.89 Å for PfRCR-Cy.003 (500,277 particles) and 3.02 Å for PfCyRPA-PfRIPR-Cy.00.03 (506,797 particles).

In both maps, the region corresponding to PfCyRPA and Cy.003 was better resolved than the rest of the complex, and PfRIPR was more poorly resolved. Therefore, particles for PfRCR-Cy.003 and PfCyRPA-PfRIPR-Cy.003 were down-sampled 2-fold and were individually subjected to 3D variability analysis in cryoSPARC^23^ with 3 orthogonal principal modes. The motion of each complex was visualised in Chimera^39^ by exporting the output of 3D variability analysis as a volume series containing 20 frames. This indicated that both complexes showed some continuous conformational heterogeneity, manifesting as a pivoting of PfRH5 and PfRIPR. This was largest in the third principal mode analysed for PfRCR-Cy.003 which showed a portion of PfRIPR to become unresolved over the volume series while the portion in contact with PfCyRPA remained mostly unchanged.

The density for PfRH5 and PfRIPR in PfRCR-Cy.003 was improved by local refinement. PfRH5 was locally refined using a soft mask around PfRH5-PfCyRPA. Before local refinement of PfRIPR, a more homogeneous subset of particles was obtained by using heterogeneous refinement and volume outputs from 3D variability analysis. This subset (253,444 particles) was signal subtracted such that it contained signal for PfRIPR only, then subject to local refinement, first using a soft mask around PfRIPR, then around a sub-portion of PfRIPR, guided by the motions observed in 3D variability analysis. The same procedure was used for PfRIPR in PfCyRPA-PfRIPR-Cy.003 except that the full particle stack for PfCyRPA-PfRIPR-Cy.003 (506,797 particles) was used.

Some 2D classes indicated that there was additional density for PfRIPR that was not resolved in these maps. To visualise this extra region, a subset of particles (62,817 particles) was processed by non-uniform refinement using the PfRCR-Cy.003 volume low-pass filtered to 30 Å as a reference. This yielded a map showing a small region of additional density projecting from the middle of the PfRIPR core, with a global resolution of 3.95 Å. Refinement with wider masks around this region did not result in more of PfRIPR becoming resolved.

The local resolution of unsharpened maps was estimated using cryoSPARC. Composite maps for PfRCR-Cy.003 and PfCyRPA-PfRIPR-Cy.003 were generated from consensus and local refinement maps in PHENIX^40^. Maps were post-processed using the default parameters of DeepEMhancer^41^ to aid with model building. All maps were rendered in ChimeraX^42^.

### Model building and refinement

To aid model building of the PfRCR-Cy.003 complex, the crystal structures of PfRH5 (PDB ID: 4U0Q, chain C)^5^ and the PfCyRPA-Cy.003 Fab complex (PDB ID: 7PI2, chains D-F)^14^, and an AlphaFold2^27^ (AlphaFold v2.1.1) predicted model of PfRIPR (20-716 except residues 484-548) were docked into the PfRCR-Cy.003 composite map as starting models using ChimeraX. Starting models were manually re-built with COOT and ISOLDE^43^ in iterative cycles. The C-terminal tail of PfRH5 and regions of PfRIPR (the N-terminus, from domain N2 to the end of EGF2, and EGFs 3 and 4) were built *de novo*, for the latter with the AlphaFold2 prediction as a guide. To build the PfCyRPA-PfRIPR-Cy.003 complex, the crystal structure of the PfCyRPA-Cy.003 Fab complex (PDB ID: 7PI2, chains D-F)^14^ and PfRIPR from the PfRCR-Cy.003 structure were rigid body fitted into the PfCyRPA-PfRIPR-Cy.003 composite map as starting models, then manually re-built using COOT. Some regions corresponding to PfRIPR were more poorly resolved in the PfCyRPA-PfRIPR-Cy.003 composite map than in the PfRCR-Cy.003 counterpart (i.e. the N-terminus and circa residues 661-667). The PfRIPR model from PfRCR-Cy.003 was unedited in these regions. Built models were refined in PHENIX using global minimisation and secondary structure restraints against their respective composite maps (which had not post-processed).

The model of PfRCR-Cy.003 comprises PfRIPR residues 34-716 (except 124-137 and 479-557), PfCyRPA residues 33-358, PfRH5 residues 159-516 (except 242-303), Cy.003 light chain residues 23-229, and Cy.003 heavy chain residues 21-245 (except 156-166). The model of PfCyRPA-PfRIPR-Cy.003 comprises PfRIPR residues 34-716 (except 124-137 and 479-558), PfCyRPA residues 34-358 (except 69-73, 124-127, 245-249 and 319-323), Cy.003 light chain residues 23-229, and Cy.003 heavy chain residues 21-245 (except 156-166).

To aid with map interpretation where density was not continuous for PfRIPR beyond Pro716, an AlphaFold2 model of PfRIPR truncated after EGF5 (residues 20-769) was generated. Docking this prediction into the PfRCR-Cy.003 map suggested that the remaining density likely corresponded to EGF5. Additionally, a model of the tail of PfRIPR (residues 717-1086, comprising EGF5 to its C-terminus) was separately predicted. This was manually docked into the second 3.95 Å PfRCR-Cy.003 map showing additional density for the tail of PfRIPR to generate a composite model of full-length PfRIPR.

To identify structural homologues of PfRIPR domains, the built structure of the PfRIPR core (residues 34-716) and the AlphaFold2 predicted structure of the PfRIPR tail (residues 717- 1086) were analysed by DALI^44^, searching against the PDB25 database.

### Microscale thermophoresis

Full-length basigin and basigin ectodomain were fluorescently labelled with Alexa Fluor™ 488 using an Alexa Fluor™ 488 protein labelling kit (Thermo Fisher) as recommended. Excess dye was removed by gel filtration on an S75 Increase 10/300 column using a buffer containing 20 mM HEPES pH 7.2, 150 mM NaCl for basigin ectodomain, or with the same buffer also containing 0.02% DDM:CHS (10:1 w/v ratio) for full-length basigin. To measure the binding of PfRH5 and PfRCR to basigin ectodomain, a two-fold dilution series of PfRH5 or PfRCR (concentration range of 8 µM to 3.91 nM) was prepared in 20 mM Tris pH 8, 200 mM NaCl, 1 mg/ml salmon sperm DNA and 0.01% Tween-20. Basigin ectodomain was held at constant 0.25 µM concentration throughout the dilution series. To measure the binding of PfRH5 and PfRCR to full-length basigin, a similar two-fold dilution series of PfRH5 and PfRCR (concentration range of 2 µM to 0.12 nM) was prepared in 20 mM Tris pH 8, 200 mM NaCl, 1 mg/ml salmon sperm DNA and 0.02% DDM. Full-length basigin was held at a constant concentration of 0.1 µM. The sample was incubated for 10 mins and centrifuged at 10,000 g for 10 mins. The supernatant was then transferred into Monolith NT.114 series premium capillaries (NanoTemper). Experiments were performed at 25 °C on a Monolith NT.115. Significant variations in raw fluorescence were observed for both PfRH5 and PfRCR measurements against full-length basigin (at >2 µM) and basigin ectodomain (at >5 µM) during data collection and so data from these concentrations were excluded from analysis. The binding experiments were performed in triplicate for two separately prepared samples. Data were analysed using software version 1.5.41 (NanoTemper).

### Surface plasmon resonance

PfRIPR and PfRIPR tail were immobilised onto separate flow paths of a CM5 Series S Sensor Chip (Cytiva) using the standard amine coupling protocol, yielding ∼8,000 response units for PfRIPR and ∼2,000 response units for PfRIPR tail. A PfCSS-PfPTRAMP complex was prepared at 8 µM by mixing each protein at a 1:1 molar ratio in SPR buffer (20 mM HEPES pH 7.5, 150 mM NaCl, 0.01% Tween-20), then a two-fold serial dilution series was prepared in the same buffer. SPR traces were recorded on a T200 Biacore instrument (Cytiva) in SPR buffer at 25 °C, with a flow rate of 30 µl/min and 180 second injections. Experiments were performed in triplicate with three independent dilution series, and binding affinities estimated by steady state analysis using T200 Biacore Evaluation Software (Cytiva).

### Crosslinking mass spectrometry of the PfRCR complex

Chemical cross-linking was performed on two separate 100 pmol aliquots (100 μl at 0.2 mg/ml) of PfRCR in 100 mM phosphate buffer pH 7.4. 2 µl of 5 mM DSSO (Thermo Fisher) in 10% DMSO, 100 mM phosphate buffer pH 7, was added for 1 hour at room temperature and quenched with 8 µl of 5% (v/v) of hydroxylamine in water (Sigma Aldrich). 1 μl of 200 mM TCEP was added for 1 hour at 55 °C and free thiol groups were alkylated with 1 μl of 380 mM Iodoacetamide (Thermo Fisher) for 30 min at room temperature in the dark. Sequential double digestions were performed with Sequencing Grade Modified Trypsin (Promega) and Sequencing Grade Chymotrypsin (Roche) at an enzyme-to-substrate weight ratio of 1:25 for 3 hours and then overnight at 37 °C in 100 mM phosphate buffer pH 7.4. Digests were diluted 1:4 with water containing 5 vol% DMSO and 0.1 vol% formic acid prior to nanoscale LC-MS analysis.

Digested protein samples were subjected to *nano*scale LC-MS analysis^45^ using a Reprosil-Pur C18-AQ trapping column (20 mm length x 100 µm I.D., 5 µm particle size, 200 Å pore size, Dr, Maisch, Ammerbuch-Entringen, Germany) and a Reprosil-Pur C18-AQ analytical column (30 cm length x 50 µm I.D., 3 µm particle size, 125 Å pore size), both packed in-house. 10 µl sample was loaded onto the trapping column at 3 µl/min of Solvent A (0.1 vol% Formic acid in water) for 10 min. The trapping column was then switched in line with the analytical column and gradients were applied from 8%-43% of solvent B (Acetonitrile + 0.1 vol% Formic acid) at 125 nl/min. The column effluent was subjected to ElectroSpray Ionization (ESI) at a spray tip voltage of 2.1 kV and a heated capillary temperature of 200°C.

Mass spectra were acquired in an Orbitrap Fusion Lumos mass spectrometer (Thermo Fisher Scientific), operated in DDA mode. Full MS scans were acquired with an Orbitrap readout (*m/z* scan range 350-1,500 Th, mass resolution of 120,000 FWHM and Normalized AGC Target of 100%). CID fragmentation spectra (employing 30% CID energy) from multiply charged (2+ to 8+) precursor ions were acquired with an Orbitrap readout at 15,000 FWHM mass resolution with the ion injection time limited to 600 msec and a Normalized AGC Target of 100%. MS3 scans, acquired to identify the peptide partners of the DSSO-cross-linked dipeptides, were triggered only upon recognition of a 31.9721-Da (±10 ppm) mass difference for a mass spectral doublet in the MS2 scan having a Partner Intensity Range for 10-100%^46^. Both doublet partners were subjected to CID fragmentation (employing 35% CID energy) at a maximum injection time of 600 msec, a Normalized AGC Target of 200% and an Ion Trap readout for the MS3 scan.

Proteome Discoverer 2.5 (Thermo Fisher Scientific) containing the XlinkX search node was used to process LC-MS data and identify crosslinked peptides. Dynamic modifications (oxidation of methionine and deamidation of asparagine) and static carbamidomethyl modification of cysteine residues were included in the search. The XlinkX search node was used with DSSO defined as MS cleavable cross-linker on lysine, serine, threonine and tyrosine residues^47^. In addition, dead-end dynamic modifications on these residues as hydrolysed or amidated DSSO were included. Mass tolerances for the MS1, MS2 and MS3 scans were set to 10 ppm, 50 ppm and 0.5 Da, respectively. The False Discovery Rate (FDR) threshold for the XlinkX Validator was set to 1%. Identified cross-links were visualised using xiNET^48^ and used to validate and aid building of the PfRCR-Cy.003 cryo-EM structure, using a Cα-Cα distance threshold of 35 Å^49^, mapped and measured using ChimeraX.

### Design and expression of cysteine-locked PfRH5

The Rosetta Disulfidize^50^ protocol was used to design cysteine locks in PfRH5. Both the wild-type and the thermally stabilised PfRH5 crystal structures (PBD ID: 6RCO^6^ and 5MI0^51^) were used for the design simulations. Following the introduction of cysteine locks, the designed models were relaxed using the Rosetta FastRelax^52^ protocol. The top scoring models based on both the total_score and dslf_fa13^53^ were manually inspected and five disulphide cysteine- lock designs connecting the N- and C-terminal halves of PfRH5 were selected for experimental validation. The Rosetta Disulfidize and Rosetta FastRelax scripts used for model design are provided in “source_data.xlsx”. The selected disulphide locks (Cys164-Cys478 (CC1), Cys180- Cys471 (CC2), Cys300-Cys408 (CC3), Cys166-Cys481 (CC4) or Cys239-Cys489 (CC5)) were introduced into PfRH5ΔNL (a recombinant construct of PfRH5 lacking its N-terminus and the α2-α3 internal loop, with the substitution C203Y (of the 7G8 *P. falciparum* strain))^5^, then wild-type PfRH5ΔNL and the cysteine-locked designs were expressed from stable S2 cell lines, as described above for full-length PfRH5. Culture supernatants were harvested after 3-4 days, 0.45 µm filtered and incubated with CaptureSelect™ C-tagXL resin (Thermo Scientific). Beads were washed using 10 column volumes of either PBS or TBS (25 mM Tris pH 7.5, 150 mM NaCl) and bound proteins eluted with 5 column volumes of C-tag elution buffer (25 mM Tris pH 7.4, 2 M MgCl2). Eluted proteins were further purified by size exclusion chromatography using a S75 Increase 10/300 column with a running buffer of PBS or TBS (25 mM Tris pH 7.5, 150 mM NaCl). Following biophysical characterisation of cysteine-lock designs CC1 to CC5, a final cysteine-locked version of PfRH5 (PfRH5^CL^) was generated through combination of cysteine-locks CC1 (Cys164-Cys478) and CC5 (Cys239-Cys489). These were introduced into PfRH5ΔNL, then expressed and purified as for other single cysteine-lock versions.

### Circular dichroism of cysteine-locked PfRH5

Circular dichroism (CD) spectra of wild-type PfRH5ΔNL and single cysteine-lock designs (CC1 to CC5) were recorded at 50 μg/ml in PBS buffer between 200 and 250 nm wavelength with a temperature ramp increasing by increments of 2 °C from 20 to 90 °C. PfRH5ΔNL and the double cysteine-locked PfRH5ΔNL^CL^ were buffer exchanged into 10 mM sodium phosphate, pH 7.5, 50 mM NaF with Zeba™ Spin Desalting Columns then CD spectra recorded at 75 μg/ml between 190 and 250 nm with temperature ramps increasing by increments of 2 °C from 20 to 90 °C. A Jasco J-815 Spectropolarimeter was used for all measurements. Data analysis was performed using the Prism 8.4.3.

### Maleimide-PEG2-biotin labelling of cysteine-locked PfRH5

PfRH5ΔNL^CL^ (10 ug) was buffer exchanged into denaturing buffer (6 M guanidine hydrochloride, 100 mM sodium acetate pH 5.5) using a 0.5 ml 7K MWCO Zeba™ Spin Desalting Column (Thermo Scientific, Cat no. 89882) and incubated at 37 degrees for 30 minutes. EZ-Link maleimide-PEG2-biotin stock (Thermo Scientific, Cat no. 21901BID) was prepared at 15 mg/ml in dimethyl sulfoxide (DMSO) and diluted to 3 mg/ml with 100 mM sodium acetate pH 5.5 immediately prior to use. Denatured PfRH5ΔNL^CL^ was labelled with a 150x molar excess of maleimide-PEG2-biotin at room temperature for 1 h. Excess maleimide-PEG2-biotin was removed by performing buffer exchange into the denaturing buffer using a 0.5 ml 7K MWCO Zeba™ Spin Desalting Column. The above steps were carried out with and without the addition of 5 mM TCEP (Thermo Scientific, Cat no. 77720) during the denaturation step. The extent of maleimide-PEG2-biotin labelling was then assessed by intact mass analysis using mass spectrometry.

### Intact mass analysis of maleimide-labelled cysteine-locked PfRH5

Reversed-phase chromatography was performed in-line prior to mass spectrometry using an Agilent 1100 HPLC system (Agilent Technologies inc. – Palo Alto, CA, USA). Concentrated protein samples were diluted to 0.02 mg/ml in 0.1% formic acid and 50 µl was injected on to a 2.1 mm x 12.5 mm Zorbax 5um 300SB-C3 guard column housed in a column oven set at 40 °C. The solvent system used consisted of 0.1% formic acid in ultra-high purity water (Millipore) (solvent A) and 0.1 % formic acid in methanol (LC-MS grade, Chromasolve) (solvent B). Chromatography was performed as follows: Initial conditions were 90 % A and 10 % B and a flow rate of 1.0 ml/min. After 15 seconds at 10 % B, a two-stage linear gradient from 10 % B to 80 % B was applied, over 45 seconds and then from 80% B to 95% B over 3 seconds. Elution then proceeded isocratically at 95 % B for 72 seconds followed by equilibration at initial conditions for a further 45 seconds. Protein intact mass was determined using a 1969 MSD-ToF electrospray ionisation orthogonal time-of-flight mass spectrometer (Agilent Technologies Inc. – Palo Alto, CA, USA). The instrument was configured with the standard ESI source and operated in positive ion mode. The ion source was operated with the capillary voltage at 4000 V, nebulizer pressure at 60 psig, drying gas at 350 °C and drying gas flow rate at 12 l/min. The instrument ion optic voltages were as follows: fragmentor 250 V, skimmer 60 V and octopole RF 250 V.

### Plasmodium falciparum culture and transfection

Blood stage *Plasmodium falciparum* parasites were cultured in human erythrocytes (UK NHS Blood and Transplant) at 3% haematocrit with custom RPMI-1640 medium supplemented with 2 mM L-glutamine, according to previously established methods^54^. All parasites used in this study were derived from a *P. falciparum* line, *p230p*DiCre, generated in strain 3D7^44^.

Parasites were synchronised by purifying schizont stages using a Percoll gradient allowing reinvasion to occur followed by sorbitol treatment of newly formed ring stages.

Transfections were performed as described^55^. Transgenic parasites were further cloned by limiting dilution. To induce DiCre recombinase mediated excision of floxed DNA, early ring stage parasites were treated with 10 nM rapamycin or DMSO as control. Parasite samples for PCR and immunoblot analysis were collected ∼40-42 hours post rapamycin treatment.

### Generation and genotyping of transgenic Plasmodium falciparum parasites

CRISPR-Cas9 guide RNA sequences targeting the 5’ region (guide 1: 5’GCTATATAAACATATTTACG-3’ and guide 2: 5’-TTTGAATTTACTATATGTAC-3’) or the 3’ region of *PfRh5* ORF (5’-TTGTCATTTCATTGTGTAAG-3’) were identified using EuPaGDT (http://grna.ctegd.uga.edu/). Each guide was cloned into vector pDC2-Cas9-hDHFRyFCU^55^, generating plasmids pDC2-Cas9-5’guide1, pDC2-Cas9-5’guide2, and pDC2-Cas9-3’guide. All primers used to generate DNA repair plasmids, along with templates and expected PCR product sizes are listed in Supplementary Tables 1 and 2.

A DNA repair template designed to replace the endogenous *PfRh5* intron with a synthetic *SERA2 LoxP* intron (‘*LoxPint*’)^26^ was synthesized with 300-350 bp homology regions (HR) spanning the *PfRh5 5’UTR* and coding regions on either side of the endogenous intron. HR1 (amplified with primers 1 and 2) started at 5’-GGTAAATGTAGGATTGTTCT-3’ and ended at 5’- ATAATGGTCAAAATTAATTT-3’, and HR2 (amplified with primers 3 and 4) spanned 5’- GATTAAGTTTTGAAAATGCA-3’ to 5’-ATCCACATTTTTATAGTCTT-3’. Both homology arms and the intervening *SERA2 LoxPint* module (amplified with primers 5/6) were stitched together using overlapping extension PCR (using primers 1 and 6, followed by primers 1 and 4). The final PCR product was inserted into pGEM-T Easy (Promega) and linearised using NcoI/SpeI and mixed with plasmid pDC2-Cas9-5’ guide1 and pDC2-Cas9-5’ guide2 before transfection, generating line PfRH5^NT^.

A DNA repair template to insert a *LoxP* sequence plus half of the *SERA2* intron (5’- <u>ATAACTTCGTATAGCATACATTATACGAAGTTAT</u>TATATATGTATATATATATATATTTATATATTTTATATTCTTTTAG; *LoxP* sequence underlined) directly after the *PfRh5* stop codon was synthesized with a 300-350 bp HR spanning the coding and *3’UTR* regions on either side of the *PfRh5* C-term Cas9 cut site. HR1 (amplified with primers 10 and 11) started at 5’- GAATTGAATATCATACAAAA and ended at 5’-GTAAGTGGTTTATTTTTTTT. HR2 (amplified with primers 12 and 13) started at 5’-AATGACAAAACATGGTATGT and ended at 5’- CAAGTACGAGCATCCGGAAC. Part of the *LoxPint* module was amplified with primers 14 and 6. All three PCR products were subsequently fused together using overlapping extension PCR with primers 10 and 6 followed by 10 and 13. The final PCR product was inserted into pGEM-T Easy (Promega), generating plasmid p*PfRh5*_C-term_*LoxP*. Finally, this plasmid was linearised with EcoRI, mixed with pDC2-Cas9-3’guide and transfected into line PfRH5^NT^, generating line PfRH5^cKO^.

To generate a parasite line containing an inducible mutant *PfRh5* gene, a second copy of *PfRh5* was designed for integration 3’ to the floxed, endogenous *PfRh5* copy. Following rapamycin induced excision of the floxed endogenous locus, the downstream *Rh5* copy would be expressed. For this, a recodonised *PfRh5* sequence was synthesised (GeneArt) starting 3’ to the endogenous intron. This recodonised sequence was then flanked by a 5’ HR (spanning the last 546bp of the endogenous PfRh5 sequence and half of *LoxPint* module described above) and a 3’ HR spanning 595bp of the *PfRh5 3’UTR*. The 5’ HR was synthesized by overlapping extension PCR (using primers 21 and 22 and 10 and 23, followed by 21 and 23), and inserted into the GeneArt generated plasmid using SacII/AflII (HR1 started at 5’- CTTTCATGTTACAATAATAA and ended at 5’-GTAAGTGGTTTATTTTTTTT). The 3’ HR was also assembled by overlapping extension PCR (using primers 13, 18, 19 and 20, followed by 18 and 20), starting at 5’-AATGACAAAACATGGTATGT and ending at 5’-TGATATAAATGAAGCGTTGA.

The final PCR product was inserted into the plasmid via MfeI/SalI, generating the final plasmid, p*PfRh5*_WT_sec_copy. Finally, the plasmid was digested using NcoI/NotI, mixed with pDC2- Cas9-3’guide and transfected into the transgenic PfRH5^NT^ parasite line, generating line PfRH5^WT^.

To generate a parasite line expressing a ‘locking cysteines’ *PfRh5* mutant upon rapamycin induced excision of the endogenous floxed *PfRh5 gene*, plasmid p*PfRh5*_WT_second_copy was modified to introduce the following mutations: L164C, E239C, M478C, and H489C. Mutations L164C and E239C were introduced with primers via overlapping extension PCR (using primers 25 and 26 and 27 and 28, followed by 25 and 29) using p*PfRh5*_WT_sec_copy as template. The final PCR product was cloned into plasmid p*PfRh5*_WT_second_copy using AflII/NdeI, yielding plasmid p*PfRh5*_lockingCyst_sec_copy_A. Likewise, mutations M478C and H489C were also introduced by PCR amplification (using primers 30 and 31 and 32 and 33, followed by 30 and 33), followed by cloning of this DNA fragment into plasmid p*PfRh5*_lockingCyst_sec_copy_A using BamHI/MfeI restriction enzymes, yielding plasmid p*PfRh5*_lockingCyst_sec_copy_B. This plasmid was digested using NcoI/NotI, mixed with pDC2-Cas9-3’guide and transfected into the transgenic PfRH5^NT^ parasite line, generating line PfRH5^CL^.

All plasmid DNA sequences were verified by Sanger sequencing. Positions of diagnostic primers used to genotype transgenic parasites are shown in schematics in Extended Data Figure 6a-b. Diagnostic primer sequences along with expected PCR product sizes are listed in Supplementary Tables 1 and 2. A positive control PCR reaction using primers 36 and 37 to amplify a 737 bp product from the *PfRON2* locus was also included in each set of diagnostic PCRs.

The Qiagen DNeasy Blood and tissue kit was used for all genomic DNA extractions. All diagnostic PCR analyses were performed using GoTaq Green (Promega) under the following conditions: 5 min at 95°C, then 33 cycles of 30 s at 95°C, 30 s at 55°C, and 1 min, 30 sec/kb at 64°C. For amplification of fragments used for construct synthesis, CloneAmp HiFi polymerase (Takara) was used. A typical reaction was run with 32 cycles of 5 s at 98°C, 15 s at 55°C, and 10 s/kb at 68°C.

### Immunoblotting

Synchronised schizonts were harvested by Percoll gradient centrifugation, washed in RPMI 1640 without albumax, and lysed in SDS sample buffer containing 100 mM DTT before protein separation on precast Bis-Tris polyacrylamide gels (MPAGE; Merck) and transferred to nitrocellulose membranes by electroblotting. Blots were blocked overnight in 5% milk in PBS with 0.2% Tween-20 and subsequently incubated with either rat anti-PfHSP70 (1/1000)^56^ or rabbit anti-PfRh5 (1/5000)^57^ followed by goat anti-rat HRP (Sigma) or Goat anti-rabbit HRP (Bio-rad). Detection by ECL was carried out using Immobilon Western Chemiluminescent HRP Substrate (Millipore).

### Parasite growth assay

To determine the growth rate of mutant parasites relative to wild-type parasites, ring stage DMSO and rapamycin-treated cultures were adjusted to a parasitaemia of ∼0.8% and a 2% haematocrit and were grown in a gassed chamber at 37°C. A starting parasitaemia was taken when parasites reached the schizont stage of the same cycle (cycle 0) and again after a further ∼40h with parasites at schizont stage in cycle 1. Parasitaemia was measured after fixation of cells with 4% paraformaldehyde, 0.1% glutaraldehyde (Sigma) in PBS for 1 h at RT, followed by incubation for 1 h at 37°C with SYBR Green I (Life Technologies).

For flow cytometry analysis, a LSR Fortessa X-20 with BD FACSDiva Software v9.0 was used with the 530/30 filter, counting 30,000 singlet events per sample. Gating for erythrocytes was achieved by plots of forward scatter area against side scatter area (gate = P1). Doublet discrimination required gating of forward scatter area against forward scatter width (gate = P2) followed by side scatter area against side scatter width (gate = P3). A SYBR Green-stained uninfected RBC sample was used as a negative control. Gating of SYBR Green positive, infected erythrocytes was achieved by plotting side scatter area against Alexa Fluor 488 area using 530/30 standard filter (gate = P4). Parasitaemia was determined by the number of cells identified in gate P4 as a percentage of those in gate P3 (Supplementary Figure 1). Data were analysed using FlowJo v10. Graphpad Prism v9.0.0 was used for statistical analysis (two-tailed, unpaired t-test) and graph generation.

### Data availability statement

Cryo-EM maps are available from the Electron Microscopy Data Bank under accession codes EMDB-16569 (PfRCR-Cy.003) and EMDB-16570 (PfCyRPA-PfRIPR-Cy.003), while coordinates are available from the Protein Data Bank under accession codes 8CDD (PfRCR-Cy.003) and 8CDE (PfCyRPA-PfRIPR-Cy.003). Uncropped gels and source data for all graphs generated in this study are provided in “source_data.xlsx”. All other data is available from the authors on request.

### Code availability statement

Rosetta scripts used in this study are provided in “source_data.xlsx”.

## Acknowledgements

This work was funded through a Wellcome Investigator award (220797/Z/20/Z) to M.K.H. Crosslinking mass spectrometry analysis of the PfRCR complex was funded through H2020 TRANSVAC2 Grant Agreement Number 730964. Cy.003 was produced through the European Commission FP7 EURIPRED project (INFRA-2012-312661), funded by the European Union’s Seventh Framework Programme [FP7/2007—2013] under Grant Agreement No: 312661— European Research Infrastructures for Poverty Related Diseases (EURIPRED). We thank the Rishi Matadeen, Joseph Caesar and Teige Matthews-Palmer at the COSMIC cryo-EM facility (University of Oxford) for support with data collection and data processing. We thank Larissa van der Maas and Hugo D. Meiring (Intravacc) and Rod Chalk (University of Oxford) for support with mass spectrometry. We would also like to thank the NHS Blood and Transplant services for the supply of human blood for parasite culture, and PlasmoDB^58^, hosted by the VEuPathDB resource centre for providing *Plasmodium* genomic DNA sequences.

## Author contributions

B.F. expressed and purified proteins, performed structure determination and surface plasmon resonance analysis. N.A. designed, produced and performed biophysical characterisation of cysteine-locked PfRH5. M.N.H. and E.K. generated transgenic parasite lines and assessed the effect of cysteine-locked PfRH5 on invasion. A.J. performed basigin binding studies. R.J.R. and S.J.D. contributed cross-linking mass spectrometry data. H.W.H contributed protein reagents.

E.K. and M.K.H. designed experiments and contributed expertise and funding. B.F. and M.K.H. prepared the manuscript and all authors contributed and commented.

## Extended Data Figures and Tables

**Extended Data Figure 1:**
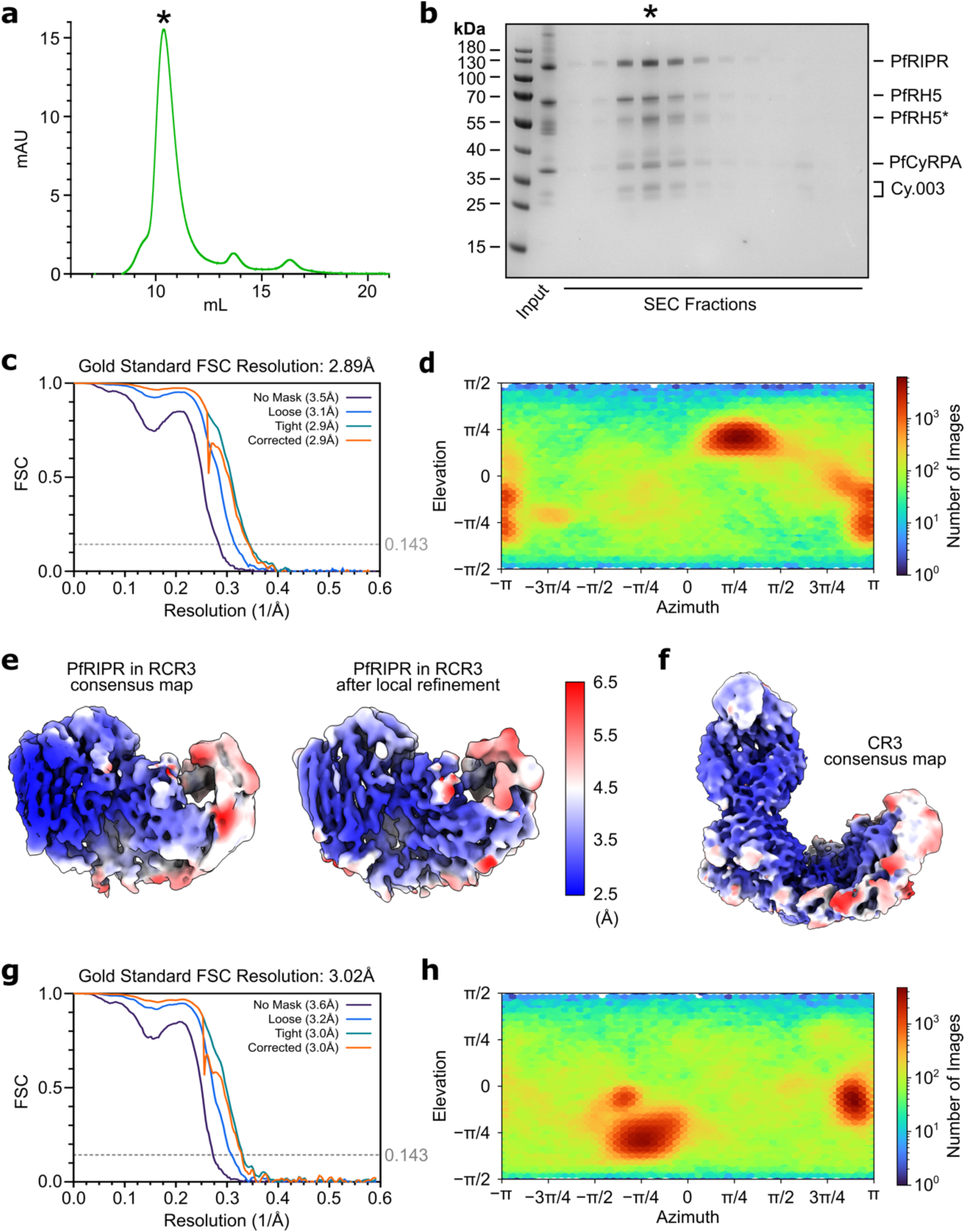
Purification of PfRCR-Cy.003 complex and cryo-EM processing statistics. **a**, Size exclusion chromatography profile and **b**, Reducing SDS-PAGE gel for the reconstituted PfRCR-Cy.003 complex. The fraction labelled with an asterisk corresponds to the region highlighted on the size exclusion chromatography profile in panel (**a**) containing the PfRCR-Cy.003 complex. Both full-length PfRH5 and truncated PfRH5 (PfRH5*) were observed in the purified complex. **c**, Gold-standard Fourier shell correlation curves of the consensus PfRCR-Cy.003 reconstruction. **d**, Particle view distribution of the consensus PfRCR-Cy.003 reconstruction. **e**, Map regions corresponding to PfRIPR in the consensus PfRCR-Cy.003 reconstruction and following local refinement, showing improvement in map quality. Both maps are coloured by local resolution. **f**, Consensus map of the PfCyRPA-PfRIPR-Cy.003 (CR3) complex coloured by local resolution as in (**e**). **g**, Gold-standard Fourier shell correlation curves of the consensus PfCyRPA-PfRIPR-Cy.003 reconstruction. **h**, Particle view distribution of the consensus PfCyRPA-PfRIPR-Cy.003 reconstruction.

**Extended Data Figure 2:**
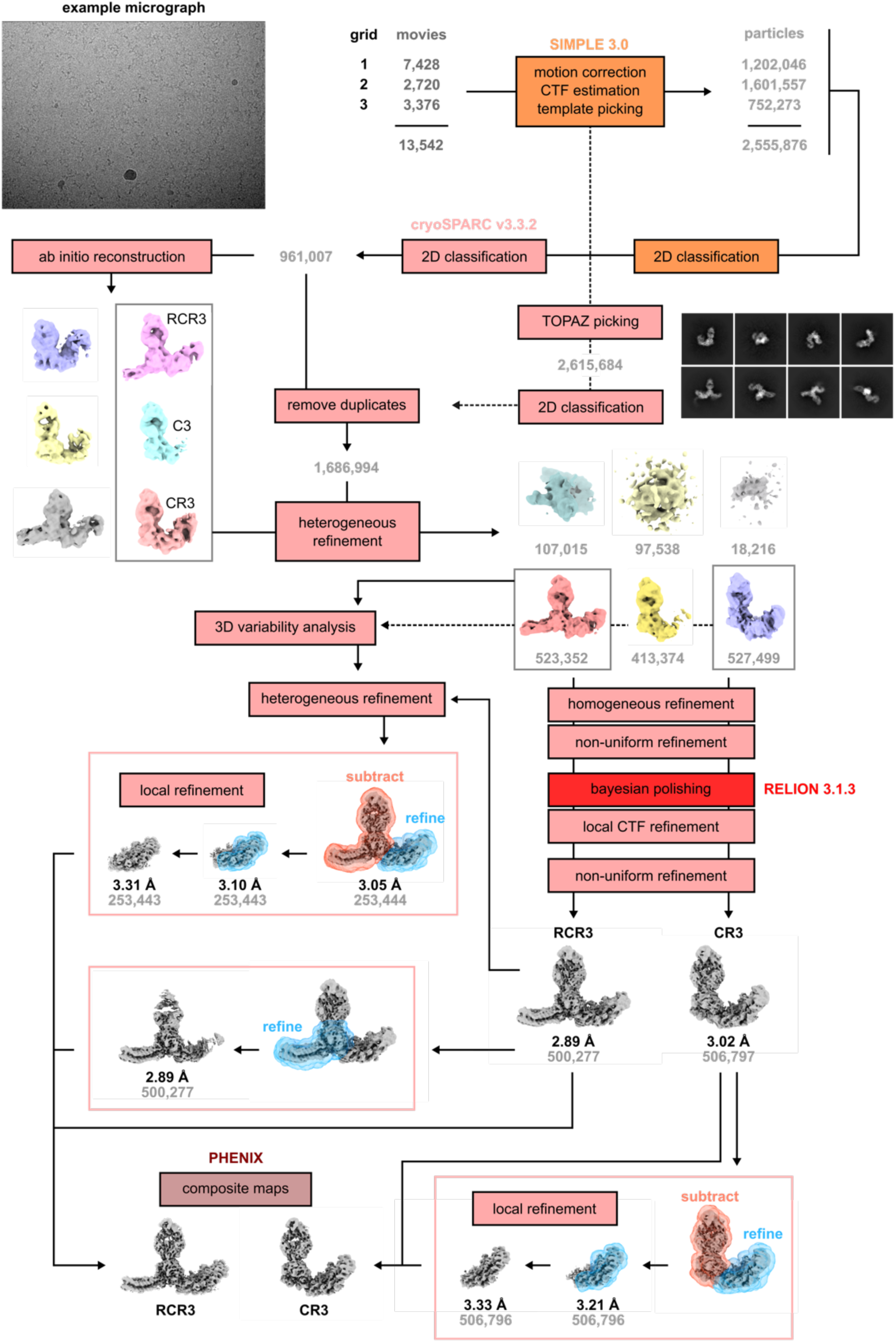
Workflow of cryo-EM data processing. Movies from three grids of the same PfRCR-Cy.003 sample were pre-processed separately in SIMPLE 3.0, followed by template-based particle picking in SIMPLE 3.0 and TOPAZ particle picking in cryoSPARC v3.3.2. After 2D classification and removal of duplicate particles, the cleaned particle stack was subject to heterogeneous refinement using maps for PfRCR-Cy.003 (RCR3), PfCyRPA-PfRIPR-Cy.003 (CR3) and PfCyRPA-Cy.003 (C3) complexes from *ab initio* reconstructions and three decoy volumes. Particle stacks corresponding to RCR3 and CR3 complexes were then separately refined in 3D space, then after Bayesian polishing in RELION 3.1.3 and local CTF refinement in cryoSPARC v3.3.2, subjected to a final non-uniform refinement to obtain consensus maps for each complex. Each map was further locally refined to improve the maps of PfRH5 and PfRIPR respectively using soft masks around the proteins of interest, and with signal subtraction for PfRIPR local refinements. Composite maps were generated using PHENIX.

**Extended Data Figure 3:**
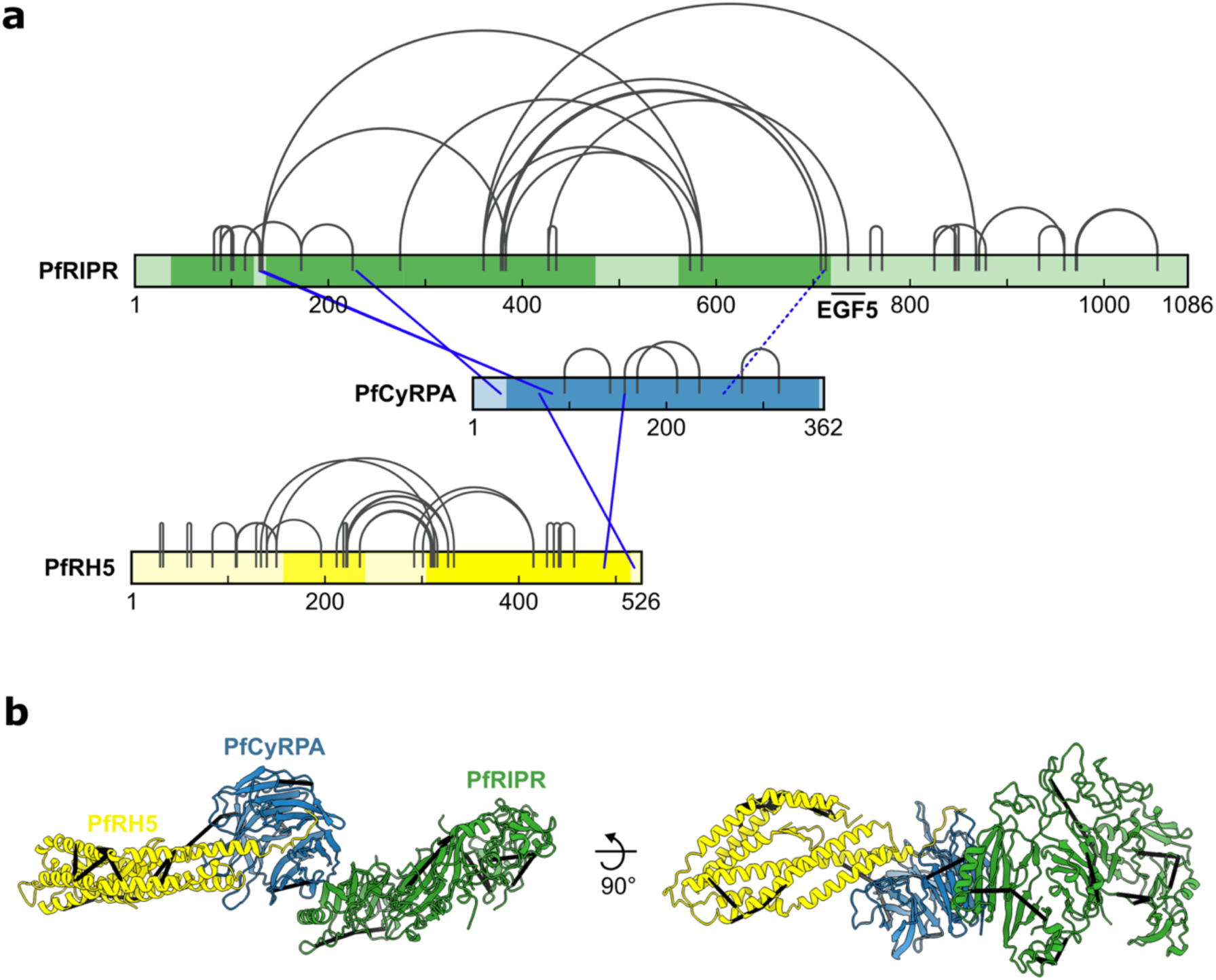
Crosslinking mass spectrometry of the PfRCR complex. **a**, Crosslinks found between PfRIPR, PfCyRPA and PfRH5 after incubating the reconstituted PfRCR complex with the crosslinker DSSO. Inter-protein crosslinks are blue and intra-protein crosslinks are grey. Crosslinks that are inconsistent with the PfRCR complex structure (with a Cα-Cα distance greater than 35Å) are indicated by dashed lines. The regions of PfRIPR, PfCyRPA and PfRH5 built in the PfRCR structure are highlighted by a darker tone. The location of EGF5 in PfRIPR is indicated to highlight the intra-PfRIPR crosslink supporting the proposed location of this domain in the PfRIPR structure. Prepared using xiNET^48^. **b**, Crosslinks with a Cα-Cα distance less than 35 Å mapped on to the PfRCR complex structure.

**Extended Data Figure 4:**
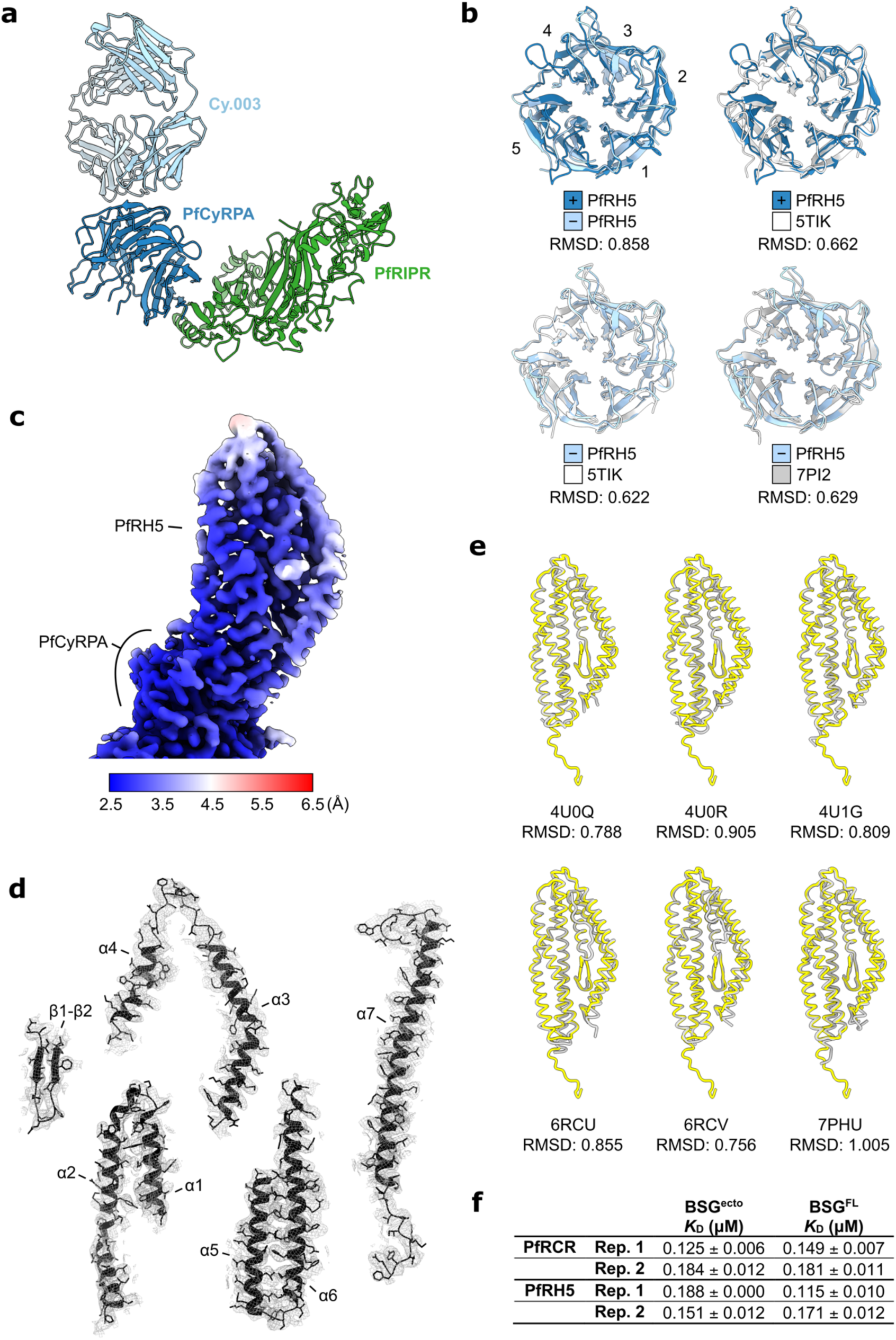
Comparison of PfRH5 and PfCyRPA structures. **a**, Structure of the PfCyRPA-PfRIPR-Cy.003 complex in cartoon representation, with PfRIPR (green), PfCyRPA (blue) and Cy.003 Fab (light blue) highlighted. **b**, Structural alignments of PfCyRPA from PfRCR-Cy.003 (dark blue), PfCyRPA-PfRIPR-Cy.003 (light blue), Cy.003-bound PfCyRPA (grey, PDB ID: 7PI2^14^), and unbound PfCyRPA (white, PDB ID: 5TIK^15^), with backbone RMSDs for each pair given below. Blades 1 through 5 are labelled. **c**, Cryo-EM map of PfRH5 following local refinement, post-processed by DeepEMhancer and coloured by local resolution. **d**, PfRH5 split into secondary structure elements, shown with their corresponding cryo-EM densities. **e**, Structural alignment of PfRH5 from PfRCR-Cy.003 (yellow) with crystal structures (grey) of basigin-bound (PDB ID: 4U0Q)^5^ or Fab-bound PfRH5 (PDB IDs: 4U0R, 4U1G, 6RCU, 6RCV, 7PHU)^5, 6^. Backbone RMSDs between PfRCR-Cy.003 and each crystal structure are given below. **f**, Binding affinity measurements for PfRCR and PfRH5 binding to basigin ectodomain (BSG^ecto^) and full-length basigin (BSG^FL^) determined by microscale thermophoresis. Values determined from two independently prepared samples are provided. Each of these was measured three times.

**Extended Data Figure 5:**
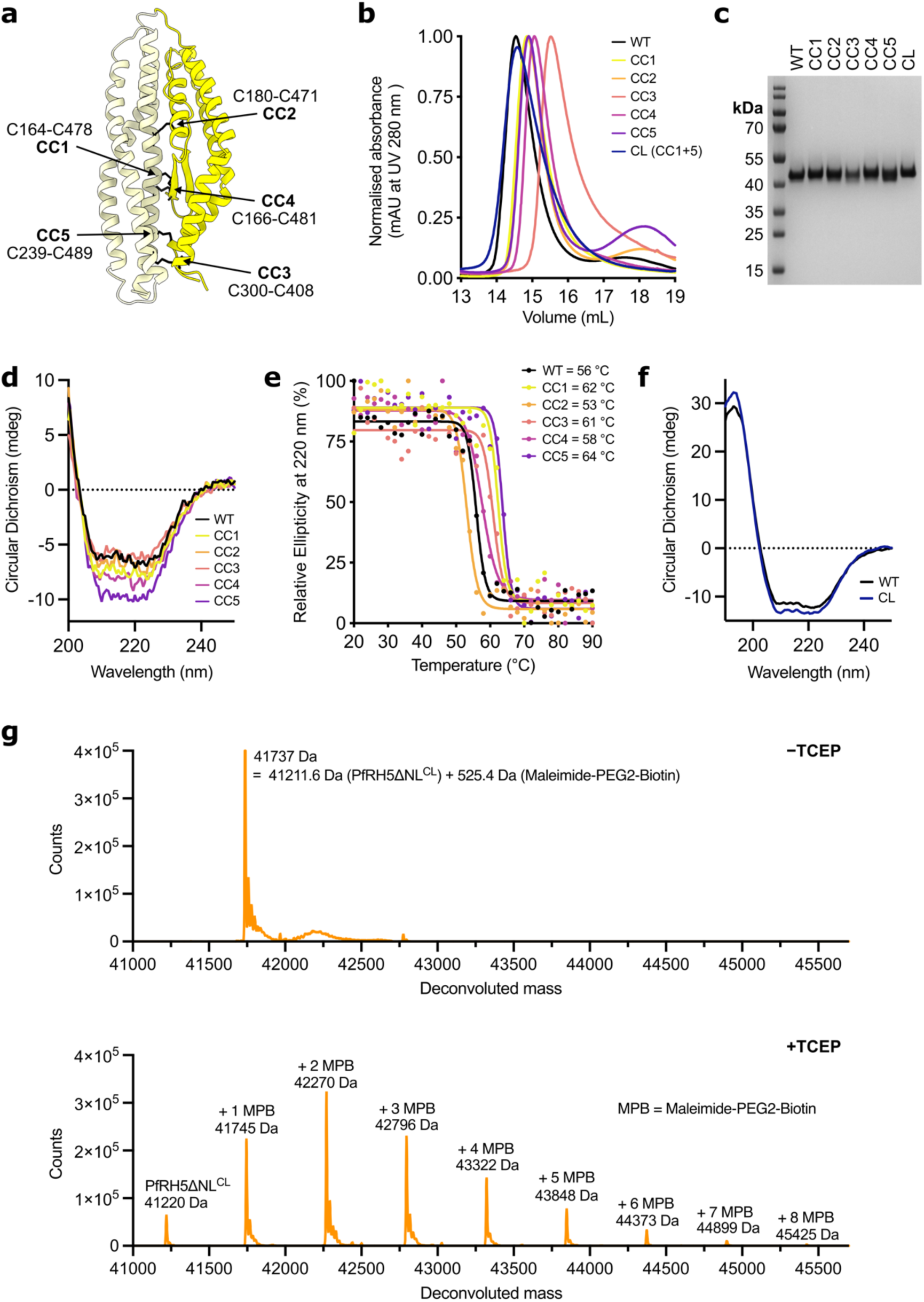
Design, expression and characterisation of cysteine-locked PfRH5. **a**, PfRH5 in cartoon showing the location of the five designed disulphide bonds (CC1 to CC5, black sticks), joining the N-terminal (dark yellow) and C-terminal (light yellow) halves of PfRH5. **b**, Gel filtration traces of purified wild-type PfRH5ΔNL (a construct lacking the unstructured N-terminus and α2-α3 loop of PfRH5^5^) and the cysteine-locked PfRH5 designs (CC1 to CC5) following expression in *Drosophila* S2 cells. **c**, SDS-PAGE gel of purified wild-type and single cysteine-locked variants (CC1 to CC5) and the double cysteine-locked (PfRH5^CL^, which combines crosslinks CC1 and CC5) version of PfRH5ΔNL. **d**, Single circular dichroism traces of wild-type and cysteine-locked variants of PfRH5ΔNL at 20 °C from their respective temperature ramps. **e**, Relative ellipticity at 220 nm versus temperature, and the calculated melting temperature (Tm) of wild-type and cysteine-locked variants of PfRH5ΔNL. **f**, Circular dichroism traces of purified wild-type and the double cysteine-locked PfRH5ΔNL (PfRH5ΔNL^CL^). **g**, Intact mass spectrometry analysis of PfRH5ΔNL^CL^ following maleimide-PEG-biotin labelling in the absence (top) and the presence of TCEP (bottom). Unlabelled PfRH5ΔNL^CL^ has a theoretical mass of 41211.6 Da and +526 Da mass differences correspond to a maleimide-PEG2-biotin addition to cysteine residues.

**Extended Data Figure 6:**
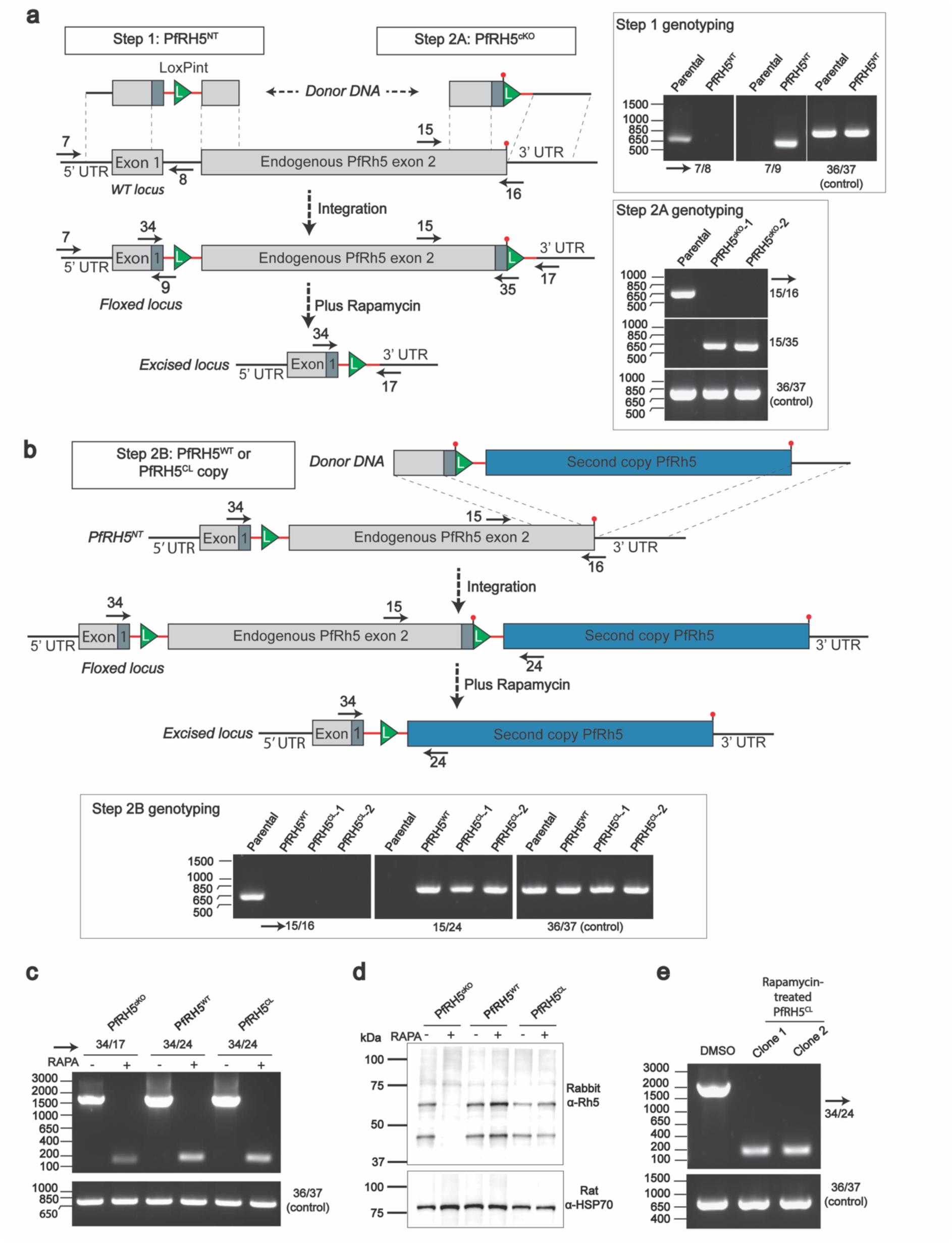
Generation of PfRH5 conditional knockout and complemented parasites. **a**, Schematic illustrating generation of transgenic rapamycin-inducible PfRH5^cKO^ line through the successive introduction of 5’ *LoxPint* (Step 1, generating line PfRH5^NT^) followed by 3’ *LoxPint* site (Step 2A), resulting in floxed exon 2 of *PfRh5* (line PfRH5^cKO^). Genotyping PCRs are shown (inset on right). One clone of PfRH5^NT^ and two clones of PfRH5^cKO^ parasites were generated. Solid black arrows indicate position of primers used to screen by PCR for integration of *LoxP* sites. Primers are numbered as in Supplementary Table 1. **b**, Schematic illustrating the introduction of a second, complementary copy of *PfRH5* (either PfRH5^WT^ or PfRH5^CL^) 3’ to endogenous *PfRH5*. The donor DNA included 548 bp of 3’ end of endogenous *PfRH5* followed by a small recodonised sequence, part of *LoxPint*, the second copy, and finally, 591 bp of endogenous 3’UTR. This design allowed for inducible complementation such that after rapamycin induced excision, only the complementary copy of *PfRH5* was expressed. PCR genotyping of Step 2B is shown in inset. One clone of PfRH5^WT^ and two clones of PfRH5^CL^ parasites were generated. For schematics in both **a** and **b**, green triangles represent *LoxP* DNA sequence; red lines represent *sera2* intron sequence; dark grey shaded boxes represent recodonised pieces of DNA. **c**, Genotyping PCR demonstrating that rapamycin induced excision of endogenous *PfRH5* exon 2. **d**, Anti-PfRH5 western blotting demonstrating ablation of PfRH5 expression for rapamycin treated PfRH5^cKO^ parasites or expression of a complementary copy of *PfRH5* for PfRH5^WT^ and PfRH5^CL^ lines. Hsp70 was used as a loading control. **e**, Rapamcyin treated PfRH5^CL^ parasites could be cloned by limiting dilution and maintained in culture. Primers used in PCR genotyping are listed in Supplementary Table 1 and the expected PCR product sizes in Supplementary Table 2. For each set of PCR reactions, control primers 36/37 were used to amplify a 737 bp product from the *PfRON2* locus.

**Extended Data Figure 7:**
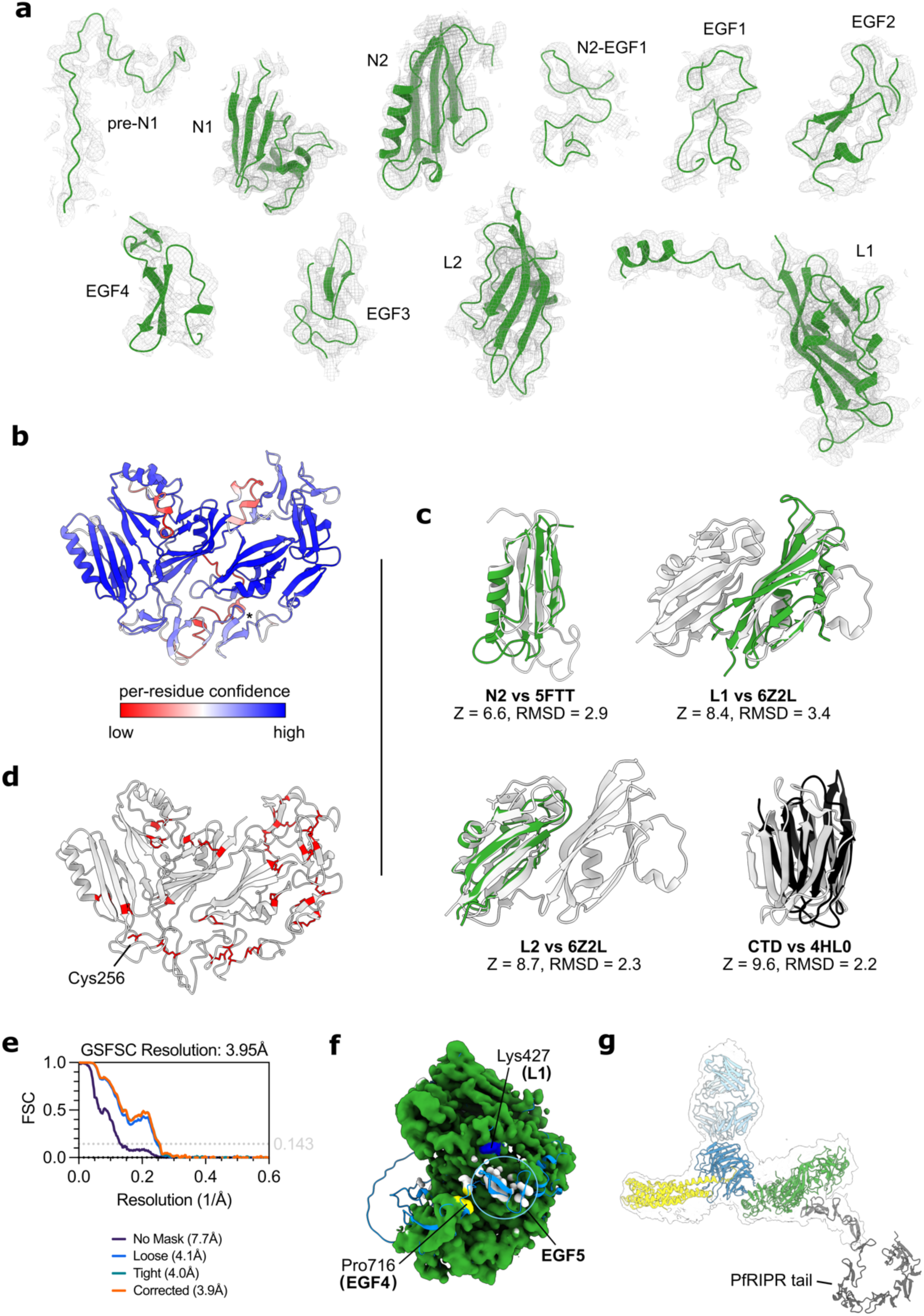
The PfRIPR core and generating a composite model of full-length PfRIPR. **a**, Domains of PfRIPR with their corresponding cryo-EM density. **b**, AlphaFold 2 prediction of PfRIPR residues 19-716 (except 484-548) coloured by pLDDT score. The regions of low confidence in this model correlate with those that required *de novo* building. **c**, Structural alignment of PfRIPR domains and their structural homologues with their DALI Z scores and RMSD scores. Structural homologues are identified by their PDB codes: 5FTT (lectin domain of Lactrophilin 3), 6Z2L (PfP113) and 4HL0 (Galectin). **d**, The PfRIPR core contains 23 disulphide bonds and one free cysteine (Cys256). Cysteine residues are shown as red sticks. **e**, Gold standard Fourier shell correlation (GSFSC) curves for the PfRCR-Cy.003 map containing additional density for the start of the PfRIPR tail. **f**, The PfRIPR portion of the composite PfRCR-Cy.003 map coloured green where a model of the PfRIPR core has been built. Uninterpretable density (coloured white) is observed in this map beyond domain EGF4 at Pro716 (yellow). An AlphaFold2 model of PfRIPR truncated after EGF5 (residues 20-769, light blue cartoon) predicts that this likely corresponds to EGF5. This is further supported by a chemical crosslink found between Lys736 of domain EGF5 and Lys427 of domain L1 (dark blue) in XL-MS analysis of the PfRCR complex (Extended Data Fig. 3). **g**, To generate a composite model of full-length PfRIPR, the AlphaFold 2 prediction of the PfRIPR tail (residues 717-1086) was docked onto the PfRIPR core using the 3.95 Å PfRCR-Cy.003 map containing additional PfRIPR density as a guide.

**Extended Data Fig 8:**
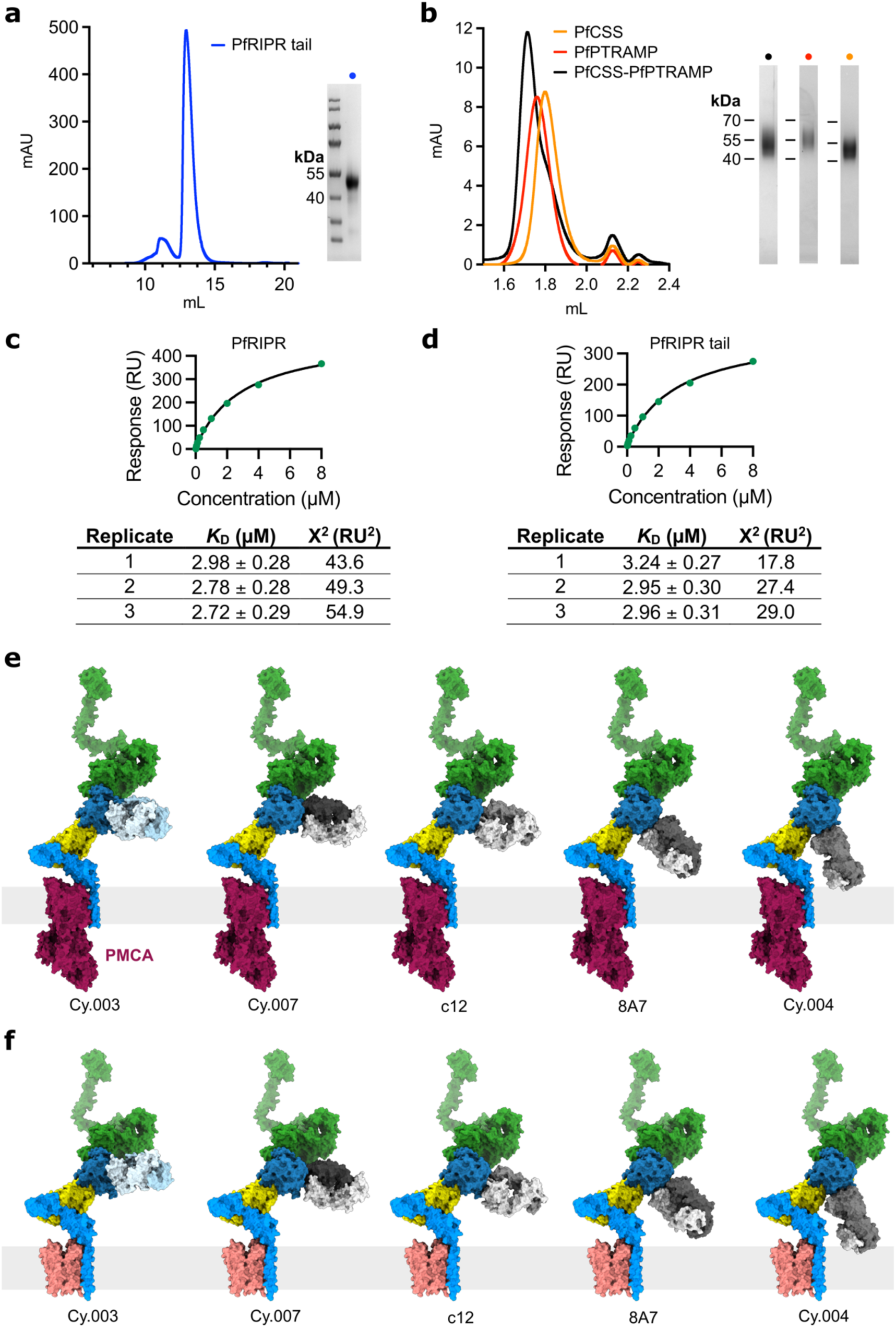
PfRIPR tail interacts with the PfCSS-PfPTRAMP complex, and the mapping of anti-PfCyRPA antibodies. **a**, Gel filtration trace of recombinantly expressed PfRIPR tail with reducing SDS-PAGE gel of isolated PfRIPR tail shown inset right. **b**, Gel filtration traces of PfCSS (orange), PfPTRAMP (red) and PfCSS-PfPTRAMP complex (black) with reducing SDS-PAGE gel inset right. SDS-PAGE fractions shown correspond to the peak fraction of each gel filtration trace. **c**, and **d**, Representative steady state fit analysis of (**c**) PfCSS-PfPTRAMP complex binding to full-length PfRIPR and (**d**) PfCSS-PfPTRAMP complex binding to PfRIPR tail by SPR. Binding analysis was completed in triplicate. Inset below are the estimated binding affinity (*K*D) and the Chi^2^ fit for each repeat. **e**, Composite model showing structural alignment of anti-PfCyRPA antibodies on to the PfRCR complex when bound to PMCA-bound basigin on the erythrocyte surface. Generated through alignment of the PfRCR-Cy.003 complex, Fab-PfCyRPA crystal structures (PDB IDs: 7PHV, 5EZO, 5TIH and 7PHW)^14–16^, and PMCA-bound basigin (PDB ID: 6A69^33^). **f**, As in (**e**) except when PfRCR is bound to MCT1-bound basigin (PDB ID: 7CKR^32^).

**Extended Data Table 1:**
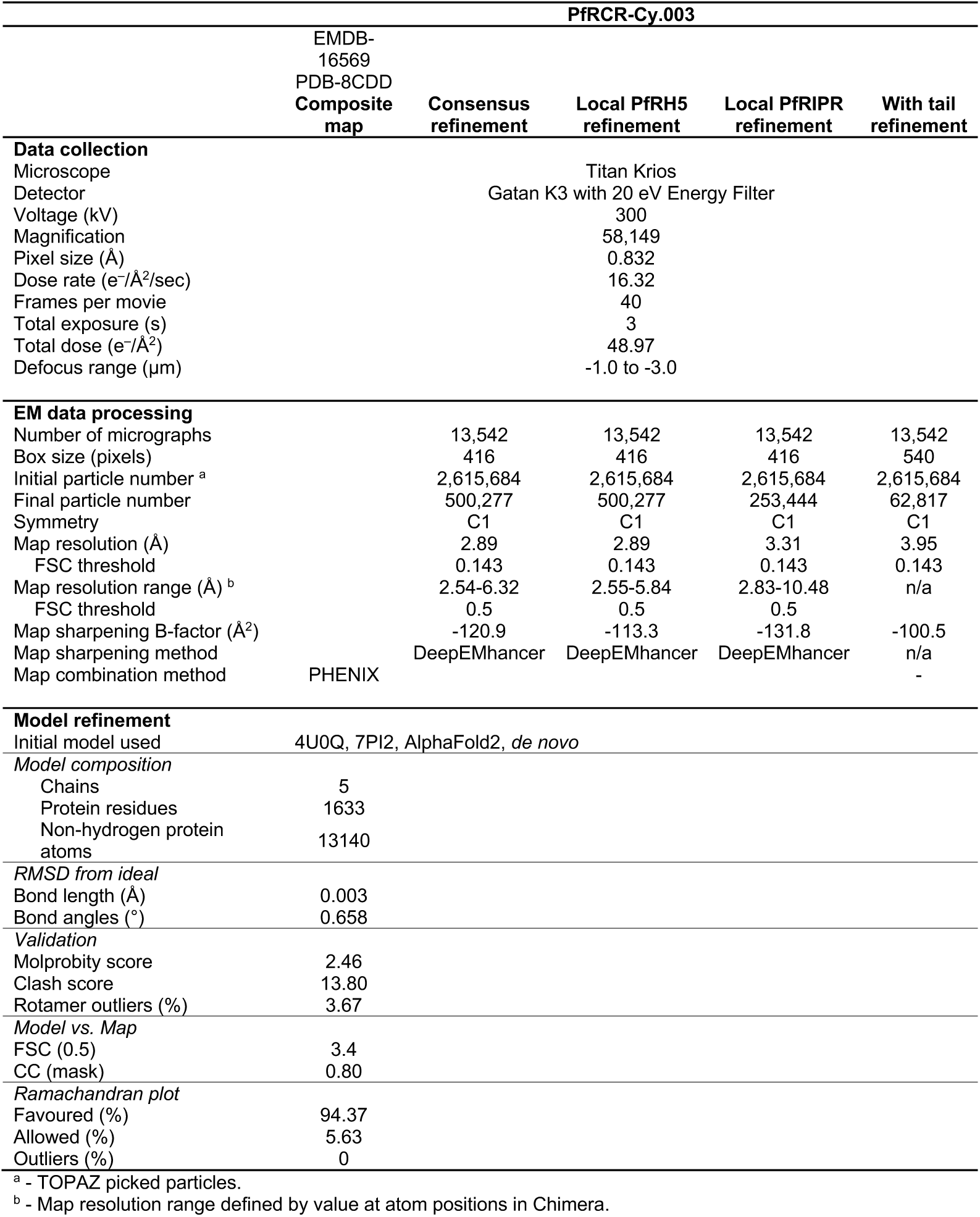
Cryo-EM data collection, refinement, and validation statistics of PfRCR-Cy.003.

| PfRCR-Cy.003 |  |  |  |  |  |
| --- | --- | --- | --- | --- | --- |
|  | EMDB-16569<br>PDB-8CDD<br>Composite map | Consensus refinement | Local PfRH5 refinement | Local PfRIPR refinement | With tail refinement |
| Data collection |  |  |  |  |  |
| Microscope |  |  | Titan Krios |  |  |
| Detector |  |  | Gatan K3 with 20 eV Energy Filter |  |  |
| Voltage (kV) |  |  | 300 |  |  |
| Magnification |  |  | 58,149 |  |  |
| Pixel size (Å) |  |  | 0.832 |  |  |
| Dose rate (e<sup>-</sup>/Å<sup>2</sup>/sec) |  |  | 16.32 |  |  |
| Frames per movie |  |  | 40 |  |  |
| Total exposure (s) |  |  | 3 |  |  |
| Total dose (e<sup>-</sup>/Å<sup>2</sup>) |  |  | 48.97 |  |  |
| Defocus range (µm) |  |  | -1.0 to -3.0 |  |  |
| EM data processing |  |  |  |  |  |
| Number of micrographs |  | 13,542 | 13,542 | 13,542 | 13,542 |
| Box size (pixels) |  | 416 | 416 | 416 | 540 |
| Initial particle number <sup>a</sup> |  | 2,615,684 | 2,615,684 | 2,615,684 | 2,615,684 |
| Final particle number |  | 500,277 | 500,277 | 253,444 | 62,817 |
| Symmetry |  | C1 | C1 | C1 | C1 |
| Map resolution (Å) |  | 2.89 | 2.89 | 3.31 | 3.95 |
| FSC threshold |  | 0.143 | 0.143 | 0.143 | 0.143 |
| Map resolution range (Å) <sup>b</sup> |  | 2.54-6.32 | 2.55-5.84 | 2.83-10.48 | n/a |
| FSC threshold |  | 0.5 | 0.5 | 0.5 |  |
| Map sharpening B-factor (Å<sup>2</sup>) |  | -120.9 | -113.3 | -131.8 | -100.5 |
| Map sharpening method |  | DeepEMhancer | DeepEMhancer | DeepEMhancer | n/a |
| Map combination method | PHENIX |  |  |  | - |
| Model refinement |  |  |  |  |  |
| Initial model used | 4U0Q, 7PI2, AlphaFold2, *de novo* |  |  |  |  |
| Model composition |  |  |  |  |  |
| Chains | 5 |  |  |  |  |
| Protein residues | 1633 |  |  |  |  |
| Non-hydrogen protein atoms | 13140 |  |  |  |  |
| RMSD from ideal |  |  |  |  |  |
| Bond length (Å) | 0.003 |  |  |  |  |
| Bond angles (°) | 0.658 |  |  |  |  |
| Validation |  |  |  |  |  |
| Molprobity score | 2.46 |  |  |  |  |
| Clash score | 13.80 |  |  |  |  |
| Rotamer outliers (%) | 3.67 |  |  |  |  |
| Model vs. Map |  |  |  |  |  |
| FSC (0.5) | 3.4 |  |  |  |  |
| CC (mask) | 0.80 |  |  |  |  |
| Ramachandran plot |  |  |  |  |  |
| Favoured (%) | 94.37 |  |  |  |  |
| Allowed (%) | 5.63 |  |  |  |  |
| Outliers (%) | 0 |  |  |  |  |

<sup>a</sup> - TOPAZ picked particles.
<sup>b</sup> - Map resolution range defined by value at atom positions in Chimera.

**Extended Data Table 2:** PfRCR mass spectrometry crosslinks. Crosslinks found between PfRIPR, PfCyRPA and PfRH5 after incubating the PfRCR complex with the crosslinker DSSO. LinkPos1 and LinkPos2 refer to the residues crosslinked from each protein. Where possible, the Cα-Cα distance measured between residues in the PfRCR structure is given.

| Type | Protein 1 | LinkPos1 | Protein 2 | LinkPos2 | Distance (Å) |
| --- | --- | --- | --- | --- | --- |
| Intra-protein | PfRIPR | 100 | PfRIPR | 89 | 12.5 |
| Intra-protein | PfRIPR | 100 | PfRIPR | 82 | 6.5 |
| Intra-protein | PfRIPR | 102 | PfRIPR | 82 | 7.2 |
| Intra-protein | PfRIPR | 129 | PfRIPR | 89 | n/a |
| Intra-protein | PfRIPR | 130 | PfRIPR | 89 | n/a |
| Intra-protein | PfRIPR | 172 | PfRIPR | 114 | 14.6 |
| Intra-protein | PfRIPR | 172 | PfRIPR | 225 | 14.3 |
| Intra-protein | PfRIPR | 274 | PfRIPR | 585 | 33.2 |
| Intra-protein | PfRIPR | 360 | PfRIPR | 573 | 14.7 |
| Intra-protein | PfRIPR | 360 | PfRIPR | 713 | 15.5 |
| Intra-protein | PfRIPR | 383 | PfRIPR | 585 | 11.8 |
| Intra-protein | PfRIPR | 383 | PfRIPR | 132 | n/a |
| Intra-protein | PfRIPR | 427 | PfRIPR | 435 | 16.4 |
| Intra-protein | PfRIPR | 585 | PfRIPR | 132 | n/a |
| Intra-protein | PfRIPR | 708 | PfRIPR | 378 | 11.8 |
| Intra-protein | PfRIPR | 708 | PfRIPR | 380 | 15.8 |
| Intra-protein | PfRIPR | 736 | PfRIPR | 427 | n/a |
| Intra-protein | PfRIPR | 759 | PfRIPR | 771 | n/a |
| Intra-protein | PfRIPR | 825 | PfRIPR | 847 | n/a |
| Intra-protein | PfRIPR | 825 | PfRIPR | 878 | n/a |
| Intra-protein | PfRIPR | 850 | PfRIPR | 846 | n/a |
| Intra-protein | PfRIPR | 868 | PfRIPR | 360 | n/a |
| Intra-protein | PfRIPR | 933 | PfRIPR | 959 | n/a |
| Intra-protein | PfRIPR | 959 | PfRIPR | 871 | n/a |
| Intra-protein | PfRIPR | 1055 | PfRIPR | 971 | n/a |
| Intra-protein | PfRIPR | 1055 | PfRIPR | 972 | n/a |
| Inter-protein | PfRIPR | 129 | PfCyRPA | 82 | n/a |
| Inter-protein | PfRIPR | 132 | PfCyRPA | 82 | n/a |
| Intra-protein | PfCyRPA | 142 | PfCyRPA | 95 | 15.2 |
| Intra-protein | PfCyRPA | 170 | PfCyRPA | 234 | 8.8 |
| Intra-protein | PfCyRPA | 211 | PfCyRPA | 157 | 13.2 |
| Intra-protein | PfCyRPA | 278 | PfCyRPA | 316 | 15.0 |
| Inter-protein | PfCyRPA | 29 | PfRIPR | 229 | n/a |
| Inter-protein | PfCyRPA | 69 | PfRH5 | 519 | n/a |
| Inter-protein | PfCyRPA | 157 | PfRH5 | 488 | 17.4 |
| Inter-protein | PfCyRPA | 259 | PfRIPR | 711 | 88.3 |
| Intra-protein | PfRH5 | 33 | PfRH5 | 30 | n/a |
| Intra-protein | PfRH5 | 58 | PfRH5 | 62 | n/a |
| Intra-protein | PfRH5 | 108 | PfRH5 | 84 | n/a |
| Intra-protein | PfRH5 | 134 | PfRH5 | 313 | n/a |
| Intra-protein | PfRH5 | 140 | PfRH5 | 129 | n/a |
| Intra-protein | PfRH5 | 150 | PfRH5 | 333 | n/a |
| Intra-protein | PfRH5 | 150 | PfRH5 | 108 | n/a |
| Intra-protein | PfRH5 | 150 | PfRH5 | 109 | n/a |
| Intra-protein | PfRH5 | 196 | PfRH5 | 140 | n/a |
| Intra-protein | PfRH5 | 219 | PfRH5 | 223 | 6.8 |
| Intra-protein | PfRH5 | 219 | PfRH5 | 222 | 5.4 |
| Intra-protein | PfRH5 | 223 | PfRH5 | 316 | 12.7 |
| Intra-protein | PfRH5 | 309 | PfRH5 | 236 | 14.5 |
| Intra-protein | PfRH5 | 311 | PfRH5 | 236 | 13.4 |
| Intra-protein | PfRH5 | 327 | PfRH5 | 223 | 14.7 |
| Intra-protein | PfRH5 | 327 | PfRH5 | 222 | 15.8 |
| Intra-protein | PfRH5 | 327 | PfRH5 | 212 | 13.9 |
| Intra-protein | PfRH5 | 415 | PfRH5 | 292 | n/a |
| Intra-protein | PfRH5 | 415 | PfRH5 | 301 | n/a |
| Intra-protein | PfRH5 | 429 | PfRH5 | 436 | 11.1 |
| Intra-protein | PfRH5 | 441 | PfRH5 | 457 | 13.3 |
| Intra-protein | PfRH5 | 443 | PfRH5 | 436 | 11.4 |

**Extended Data Table 3:** Cryo-EM data collection, refinement, and validation statistics for PfCyRPA-PfRIPR-Cy.003.

| PfCyRPA-PfRIPR-Cy.003 |  |  |  |
| --- | --- | --- | --- |
|  | EMDB-16570<br>PDB-8CDE<br>Composite<br>map | Consensus<br>refinement | Local PfRIPR<br>refinement |
| Data collection |  |  |  |
| Microscope |  | Titan Krios |  |
| Detector |  | Gatan K3 with 20 eV Energy Filter |  |
| Voltage (kV) |  | 300 |  |
| Magnification |  | 58,149 |  |
| Pixel size (Å) |  | 0.832 |  |
| Dose rate (e<sup>-</sup>/Å<sup>2</sup>/sec) |  | 16.32 |  |
| Frames per movie |  | 40 |  |
| Total exposure (s) |  | 3 |  |
| Total dose (e<sup>-</sup>/Å<sup>2</sup>) |  | 48.97 |  |
| Defocus range (µm) |  | -1.0 to -3.0 |  |
| EM data processing |  |  |  |
| Number of micrographs |  | 13,542 | 13,542 |
| Box size (pixels) |  | 416 | 416 |
| Initial particle number <sup>a</sup> |  | 2,615,684 | 2,615,684 |
| Final particle number |  | 506,797 | 506,796 |
| Symmetry |  | C1 | C1 |
| Map resolution (Å) |  | 3.02 | 3.33 |
| FSC threshold |  | 0.143 | 0.143 |
| Map resolution range (Å) <sup>b</sup> |  | 2.61-6.90 | 2.55-8.78 |
| FSC threshold |  | 0.5 | 0.5 |
| Map sharpening B-factor (Å<sup>2</sup>) |  | -131.1 | -141.2 |
| Map sharpening method |  | DeepEMhancer | DeepEMhancer |
| Map combination method | PHENIX |  |  |
| Model refinement |  |  |  |
| Initial model used | PfRCR-Cy.003 |  |  |
| Model composition |  |  |  |
| Chains | 4 |  |  |
| Protein residues | 1317 |  |  |
| Non-hydrogen protein atoms | 10465 |  |  |
| RMSD from ideal |  |  |  |
| Bond length (Å) | 0.003 |  |  |
| Bond angles (°) | 0.713 |  |  |
| Validation |  |  |  |
| Molprobit score | 2.38 |  |  |
| Clash score | 8.14 |  |  |
| Rotamer outliers (%) | 4.02 |  |  |
| Model vs. Map |  |  |  |
| FSC (0.5) | 3.6 |  |  |
| CC (mask) | 0.74 |  |  |
| Ramachandran plot |  |  |  |
| Favoured (%) | 92.12 |  |  |
| Allowed (%) | 7.88 |  |  |
| Outliers (%) | 0 |  |  |

<sup>a</sup> - TOPAZ picked particles.
<sup>b</sup> - Map resolution range defined by value at atom positions in Chimera.

**Extended Data Table 4:**
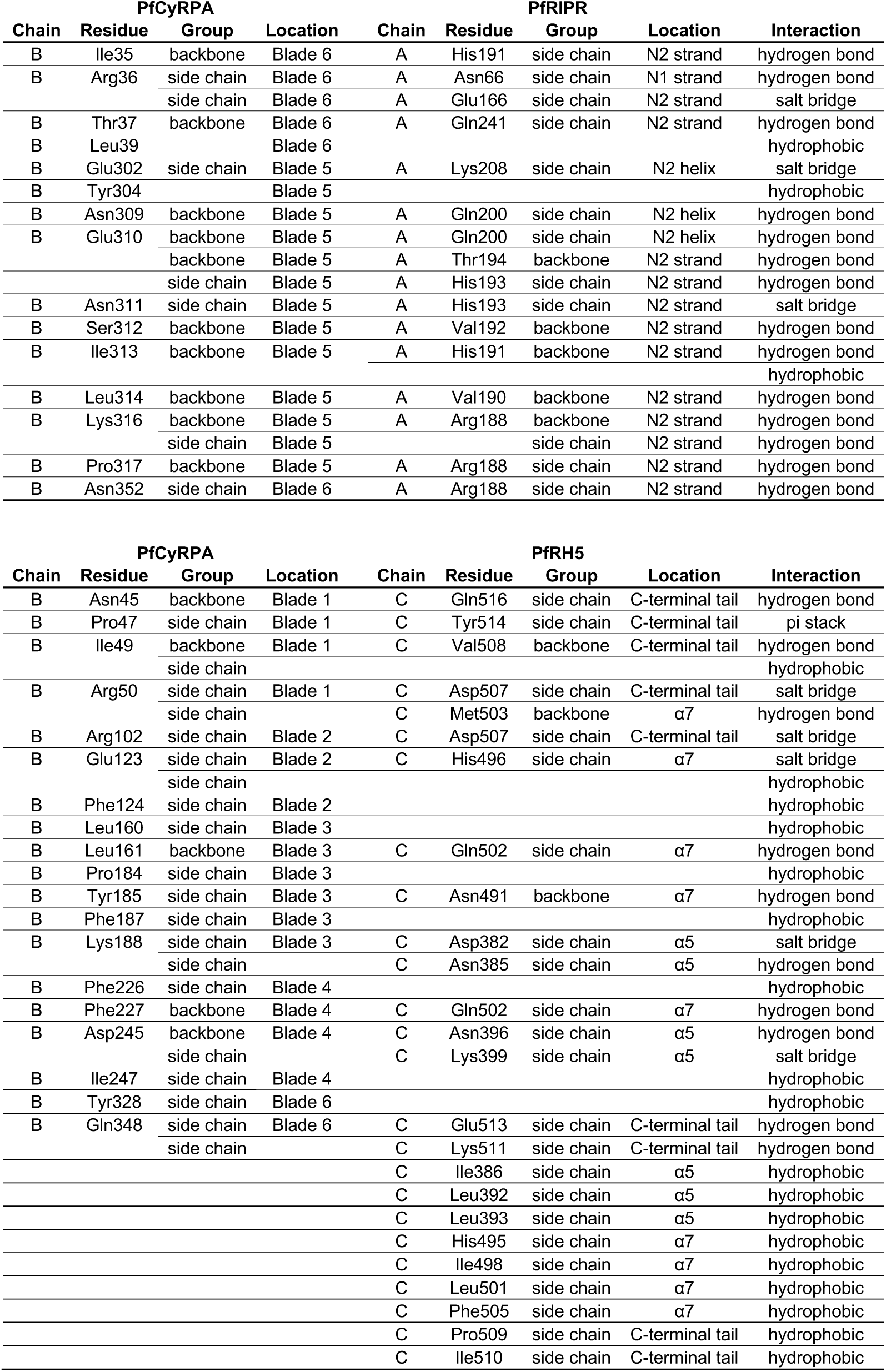
Interface residues of the PfRCR complex.

**Supplementary Table 1:**
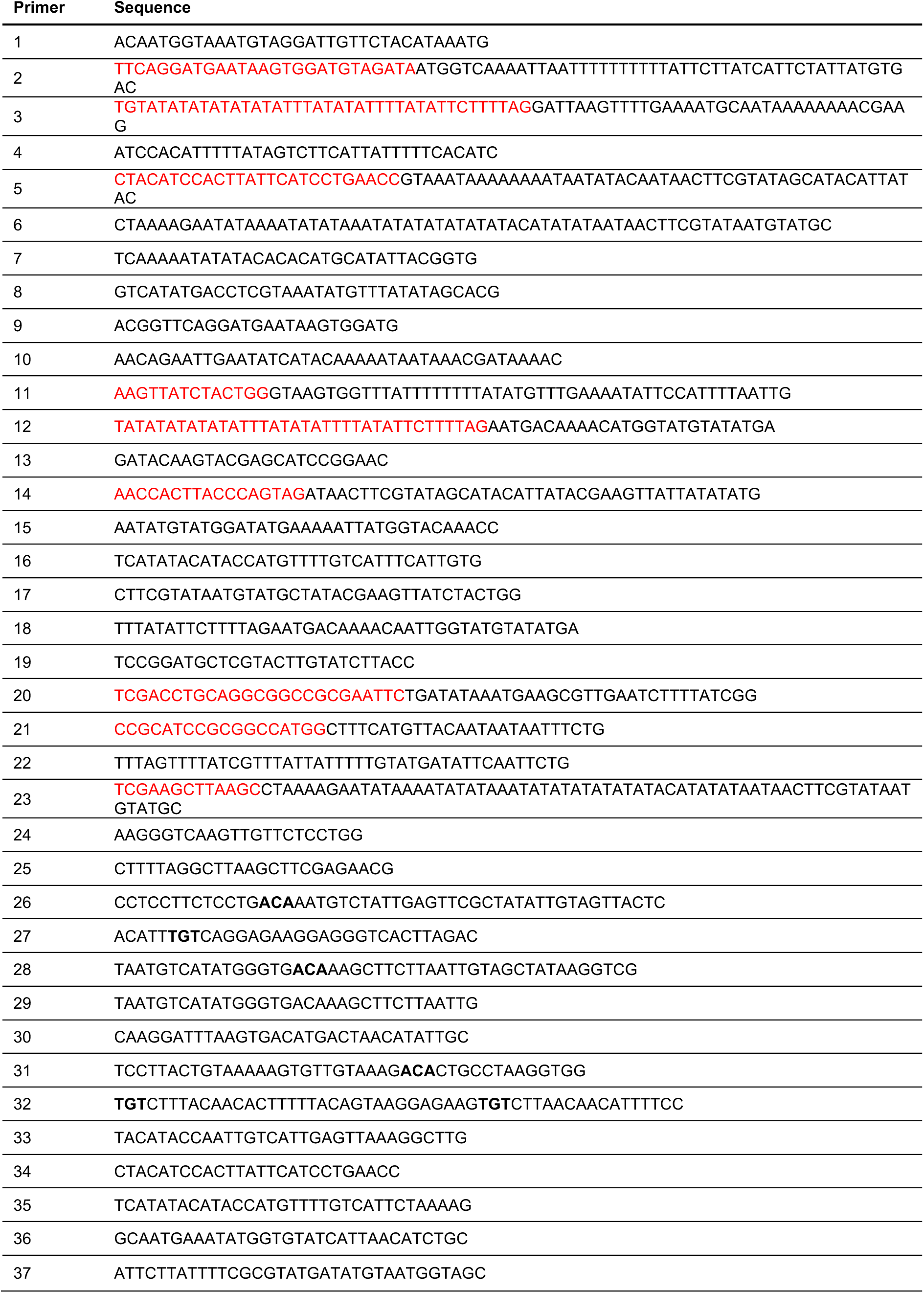
Primer numbers and sequences used in this study. Red text is overhangs introduced to PCR products for stitching/restriction cloning purposes. Bold text is sequences to introduce locking cysteine mutations into PCR products.

**Supplementary Table 2:** PCR reactions used for synthesising homologues repair donor DNA plasmids and genotyping transgenic parasites. White boxes indicate PCR reactions used for construct generation. Green boxes indicate genotyping/diagnostic PCRs.

| PCR Product / target locus | Fwd primer | Rev primer | Product size (bp) | Template(s) |
| --- | --- | --- | --- | --- |
| Generation of *PfRH5* N-term *LoxP* donor DNA plasmid (white)/ genotyping of transgenic line (green) |  |  |  |  |
| HR1 | 1 | 2 | 343 | Pf 3D7 gDNA |
| HR2 | 3 | 4 | 390 | Pf 3D7 gDNA |
| LoxPint | 5 | 6 | 129 | SERA2 LoxPint |
| HR1-LoxPint fusion | 1 | 6 | 448 | HR1 + LoxPint PCR products |
| Full length insert | 1 | 4 | 798 | HR1-LoxPint + HR2 PCR products |
| 'WT' locus | 7 | 8 | 675 | gDNA from transfection |
| Integrated locus | 7 | 9 | 585 | gDNA from transfection |
| Generation of *PfRH5* C-term *LoxP* donor DNA plasmid (white)/ genotyping of transgenic line (green) |  |  |  |  |
| HR1 | 10 | 11 | 325 | Pf 3D7 gDNA |
| HR2 | 12 | 13 | 390 | Pf 3D7 gDNA |
| LoxPint | 14 | 6 | 97 | SERA2 LoxPint |
| HR1-LoxPint fusion | 10 | 6 | 398 | HR1 + LoxPint PCR products |
| Full length insert | 10 | 13 | 752 | HR1-LoxPint + HR2 PCR products |
| 'WT' locus | 15 | 16 | 627 | gDNA from transfection |
| Integrated locus | 15 | 17 | 631 | gDNA from transfection |
| Generation of *PfRH5* WT second copy donor DNA plasmid (white)/ genotyping of transgenic line (green) |  |  |  |  |
| HR2 fragment 1 | 18 | 13 | 371 | PfRh5 C-term *LoxP* plasmid |
| HR2 fragment 2 | 19 | 20 | 296 | Pf 3D7 gDNA |
| HR2 full length | 18 | 20 | 645 | HR2.1 + HR2.2 PCR products |
| HR1 fragment 1 | 21 | 22 | 300 | PfRh5 C-term *LoxP* plasmid |
| HR1 fragment 2 | 10 | 23 | 411 | Pf 3D7 gDNA |
| HR1 full length | 21 | 23 | 666 | HR1.1 + HR1.2 PCR products |
| WT locus | 15 | 16 | 627 | gDNA from transfection |
| Integrated locus | 15 | 24 | 740 | gDNA from transfection |

| Generation of *PfRh5* locking cysteines second copy donor DNA plasmid (white)<br>/ genotyping of transgenic line (green) |  |  |  |  |
| --- | --- | --- | --- | --- |
| Fragment 1.1 | 25 | 26 | 448 | PfRH5 WT second copy plasmid |
| Fragment 1.2 | 27 | 28 | 249 | PfRH5 WT second copy plasmid |
| Fragment 1 full length | 25 | 29 | 676 | Fragments 1.1 + 1.2 |
| Fragment 2.1 | 30 | 31 | 330 | PfRH5 WT second copy plasmid |
| Fragment 2.2 | 32 | 33 | 163 | PfRH5 WT second copy plasmid |
| Fragment 2 full length | 30 | 33 | 464 | Fragments 2.1 + 2.2 |
| WT locus | 15 | 16 | 627 | gDNA from transfection |
| Integrated locus | 15 | 24 | 740 | gDNA from transfection |
| Diagnostic PCRs to detect rapamycin induced excision of floxed *PfRh5* |  |  |  |  |
| PfRH5 cKO excised/non excised locus | 34 | 35 | 1752 if non-excised; 155 if excised | gDNA from Rap/DMSO treated parasites |
| PfRH5 cKO + WT second copy excised/non excised locus | 34 | 24 | 1785 if non-excised; 188 if excised | gDNA from Rap/DMSO treated parasites |
| PfRH5 cKO + locking cysteines second copy excised/non excised locus | 34 | 24 | 1785 if non-excised; 188 if excised | gDNA from Rap/DMSO treated parasites |
| Positive control for diagnostic PCRs |  |  |  |  |
| PfRON2 locus (positive control) | 36 | 37 | 737 | gDNA from transgenic parasites |

**Supplementary Figure 1:**
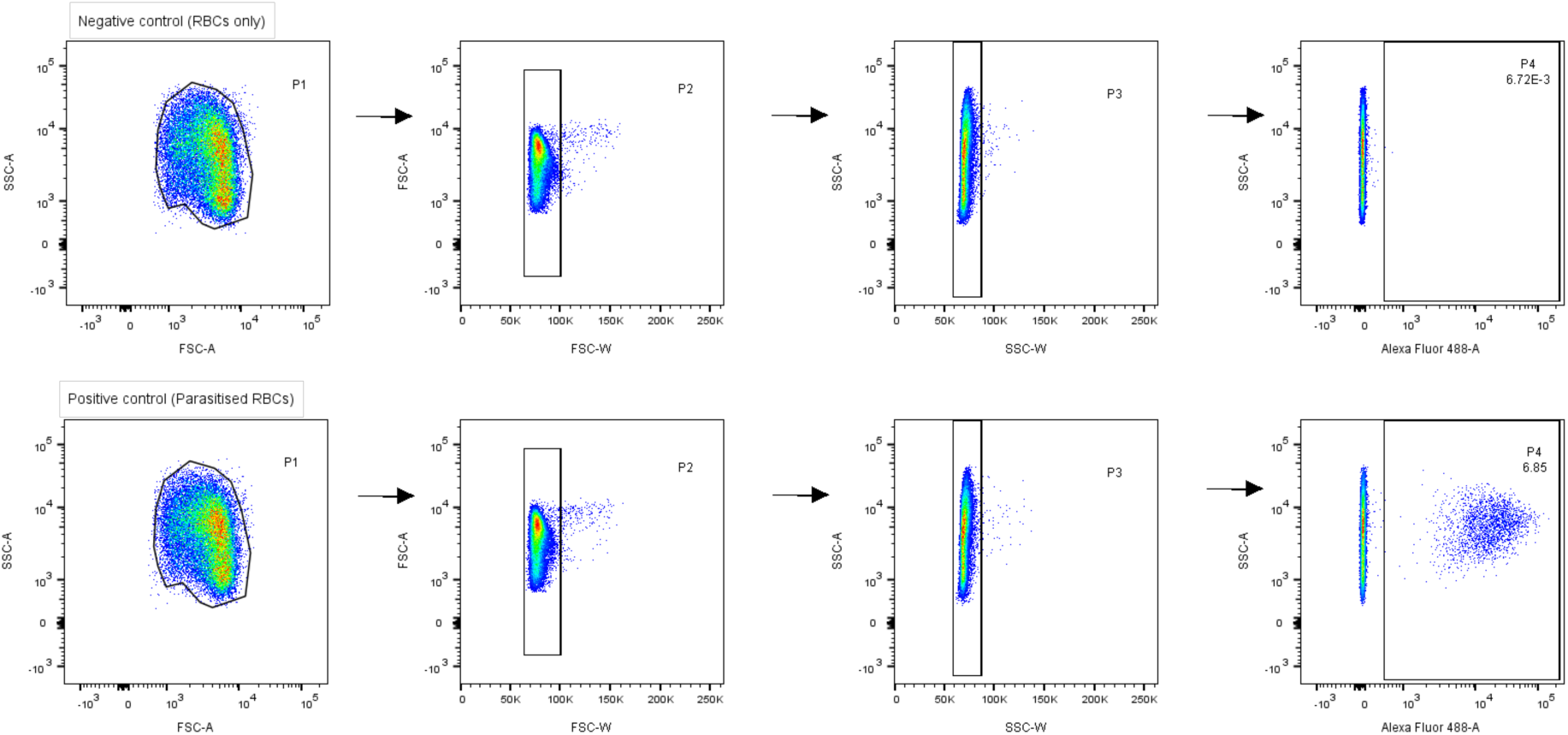
Flow cytometry gating strategies used to analyse the proportion of SYBR-positive, parasite infected erythrocytes vs uninfected erythrocytes within a given sample. Top panels indicate an erythrocyte (RBC) only, negative control. Bottom panels depict a representative sample containing parasitised erythrocytes. For each set of panels, the percentage of SYBR positive cells is indicated under gate P4. Staining and gating strategies are described in full in the materials and methods section.

